# Evolution of the recombination regulator PRDM9 in minke whales

**DOI:** 10.1101/2020.12.11.422147

**Authors:** Elena Damm, Kristian K Ullrich, William B Amos, Linda Odenthal-Hesse

## Abstract

We explored the structure and variability of the *Prdm9* gene, which codes for the PRDM9 protein, in samples of the minke whales from the Atlantic, Pacific and Southern Oceans. The PRDM9 protein controls the reshuffling of parental genomes in most metazoans and we show that minke whale possess all the features characteristic of PRDM9-directed recombination initiation, including complete KRAB, SSXRD and SET domains and a rapidly evolving array of C2H2-type-Zincfingers (ZnF). We uncovered eighteen novel PRDM9 variants and evidence of rapid evolution, particularly at DNA-recognizing positions that evolve under positive selection. At different geographical scales, we observed extensive *Prdm9* allelic diversity in Antarctic minke whales (*Balaenoptera bonarensis*) that, conversely, lack observable population differentiation in mitochondrial DNA and microsatellites. In contrast, a single PRDM9 variant is shared between all Common Minke whales and even across subspecies boundaries of North Atlantic (*B. a. acutorostrata*) and North Pacific (*B. a. scammoni*) minke whale, which do show clear population differentiation. PRDM9 variation of whales predicts distinct recombination initiation landscapes genome-wide, which has possible consequences for speciation.

## Introduction

The gene *Prdm9* encodes “PR-domain-containing 9” (PRDM9), a meiosis-specific four-domain protein that regulates meiotic recombination in mammalian genomes. The four functional domains of the PRDM9 protein are essential for the placement of double-stranded DNA breaks (DSBs) at sequence-specific target sites. Three of the domains are highly conserved: *i)* the N-terminal Kruppel-associated box- domain (KRAB) that promotes protein-protein binding, for example with EWSR, CXXC1, CDYL and EHMT2 [1, 2]; *ii)* the SSX-repression-domain (SSXRD) of yet unknown function; *iii)* the central PR/SET domain, a subclass of the SET domain, with methyltransferase activity at H3K4me3 and H3K36me3. The fourth, C-terminal, domain comprises an array of type Cystin_2_Histidin_2_ zinc-fingers (C_2_H_2_-ZnF), encoded by a minisatellite-like sequence of 84 bp tandem repeats. This coding minisatellite reveals evidence of positive selection and concerted evolution, with many functional variants having been found in humans [3, 4], mice [5, 6], non-human primates [7] and other mammals [8, 9]. Even highly domesticated species like equids [8], bovids [10] and ruminants [11, 12] show high diversity and rapid evolution. In light of this extreme variability between minisatellite-like repeat units in most other mammals, the allele found in the genome of North Pacific minke whale (*Balaenoptera acutorostrata scammoni)* is puzzling, as it comprises of an array of only a single type of C_2_H_2_-ZnF type repeat unit.

Minke whales are marine mammals of the genus *Balaenoptera*, in the order of baleen whales (*Cetaceans*), that are of particular interest not only because little is known about their population biology, seasonal migration routes and breeding behavior, but also to support future conservation efforts. Minke whales were long considered to be a single species, but have been classified as two distinct species, the Common minke whale (*Balaenoptera acutorostrata)* and the Antarctic minke whale (*Balaenoptera bonaerensis* Burmeister, 1867). The Common minke whale (*B. acutorostrata)* is cosmopolitan in the waters of the Northern Hemisphere. And can be separated into two subspecies, the Atlantic minke whale *(B. acutorostrata acutorostrata)* and the North Pacific minke whale (*B. acutorostrata scammoni*), separated from each other by landmasses and the polar ice cap. Antarctic minke whale*s* inhabit the waters of the Antarctic ocean in the Southern Hemisphere during feeding season but seasonally migrate to the temperate waters near the Equator during the breeding season [13].

Although overlapping habitats exist near the Equator, seasonal differences in migration and breeding behavior largely prevent inter-breeding between Common and Antarctic minke whales [14]. Despite this, occasional migration across the Equator has been observed [15] and recent studies have uncovered two instances of hybrid individuals, both females and one with a calf most likely sired by an Antarctic minke whale [16, 17]. Thus, hybrids may be both viable and fertile, allowing backcrossing [17]. However, it is unclear whether occasional hybridization events have always occurred or whether they are a recent phenomenon driven by anthropogenic changes, including climate change [17]. More importantly, since both hybrids were female, current data do not exclude postzygotic reproductive isolation through males, as expected under Haldane’s rule. Hybrid sterility is a universal phenomenon observed in many eukaryotic inter-species hybrids, including yeast, plants, insects, birds, and mammals [18, 19]. Minke whales offer an unusual opportunity to study *Prdm9*, the only hybrid sterility gene that is known in vertebrates thus far, sometimes called a speciation gene [20].

## Material and Methods

### PRDM9 occurrence and protein domain prediction in diverse taxa

To infer PRDM9 occurrence and to subsequent investigate protein domain architecture in diverse taxa, the genome sequences were downloaded as given in **Supplementary Table 2**. To infer PRDM9 occurrence in the given species, the annotated protein XP_028019884.1 from *Balaenoptera acutorostrata scammoni* was used as the query protein with exonerate [21] (v2.2.0) and the --protein2genome model to extract the best hit. Subsequently, for each species, the extracted coding sequences (CDS) were translated and investigated with InterProScan [22] (v5.46-81.0) and HMMER3 [23] (v3.3) using KRAB, SSXRD, SET and C_2_H_2_-ZnF as bait to obtain the protein domain architecture.

### Phylogenetic Analyses

For the phylogenetic reconstruction across Cetartiodactyla, we used only KRAB, SSXRD and SET domains (Supplementary Figure 1). These protein domains were concatenated and used as input for the software BAli-phy [24] (v3.5.0). BAli-phy was run twice with 10,000 iterations each and the default settings for amino acid input. Subsequently, the majority consensus tree was obtained by skipping the first 10% of trees as burn-in, rooted on the branch outside the Cetacea and visualized in Figtree (http://tree.bio.ed.ac.uk/software/figtree/; v1.4.4).

The phylogenetic tree from the mitochondrial D-loop region was reconstructed, including several whale species as given in Supplementary Table 3. The corresponding D-loop was aligned with MAFFT version 5 [25](v7.471) with the L-INS-i algorithm and manually curated. The maximum-likelihood tree was calculated under the TN+F+I+G4 model using IQ-TREE [26] (v1.6.12) mid-point rooted and visualised in Figtree (http://tree.bio.ed.ac.uk/software/figtree/; v1.4.4).

### Long-range sequencing of the Prdm9 gene

We first extracted the full-length DNA sequence of the PRDM9 Protein from the minke whale genome from UCSC genome browser and designed primers to flank the entire protein-coding portion, encompassing the KRAB, SSXDR, PR/SET and C_2_H_2_-ZnF binding domains using Primer-BLAST [27](Supplementary Table 4). We amplified the *Balaenoptera acutorostrata Prdm9* gene, including all introns and exons across a ~11kb interval by long-range PCR. We then sequenced the entire interval using phased long-range Nanopore Sequencing with MinION (Oxford Nanopore). Whole-length consensus sequences generated from 639 sequencing reads that cover the entire 10582 bp, with an average per base pair coverage of 269x. We used this consensus sequence as reference for *in-silico* predictions of functional domains and mapped human PRDM9 Exons 3 – 11 from ENSEMBL (ENST00000296682.3). By manually splicing all unaligned sequence fragments, we generated an in-silico predicted mRNA of *Balaenoptera acutorostrata* PRDM9. This sequence was then submitted as ‘.fasta’ to the Entrez Conserved Domains Database (CDD) home page, (https://www.ncbi.nlm.nih.gov/Structure/cdd/wrpsb.cgi) with the following parameters; searching against Database CDD v3.18 – 52910 PSSMs Expect Value threshold: 0,01, without low-complexity filter, composition-based statistics adjustment, rescuing borderline hits (ON) a maximum number of 500 hits and concise result mode).

### Wild minke whale samples

Performed investigations are not considered to be animal trials in accordance with the German animal welfare act, since samples were obtained from commercial whaling between 1980 and 1984. A total of n=143 DNA samples of minke whale with vague information about the subspecies, from four different defined commercial whaling areas. These include North Atlantic (NA; n=17) and North Pacific (NP; n=4) individuals captured during migration season, as well as Antarctic Ocean Areas IV (AN IV; n=65) and V (AN V; n=57) were obtained during feeding season. A subset of these samples had been used in van Pijlen et al., 1995 [28]. Samples from the Antarctic areas included numerous duplicate samples that were used as internal controls, analyzed for all parameters and excluded from the sample set once identical results had been confirmed (318, 325, 327, 1661, 1663).

For the detailed DNA-extraction protocol, precise catch data, as well as providers of the samples of most individuals used in this project, see van Pijlen, 1994. Briefly, genomic DNA was extracted from the skin (NA) and muscle biopsies (NP, AN) with Phenol-Chlorophorm and stored in TE at −80°C in the 1980s. If the DNA was dried out it was first eluted with TE. All DNA concentrations were determined with fluorescent Nanodrop-1000 (Thermo Fisher Scientific). For all performed analysis, 20 ng/μl DNA working stocks were prepared and stored at −20°C.

### STR genotyping

Nine autosomal, as well as X/Y microsatellite loci with di- or tetramer repeat motifs, were analysed for all samples: EV001, EV037 [29], GATA028, GATA098, GATA417 [30], GT023, GT310, GT509, GT575 [31] and sex loci X and Y [32]. Four separate multiplexing reactions were performed for each individual, and each contained 40 ng of DNA, 0.2 μM of each primer, 5 mM Multiplex-Kit (Qiagen) and HPLC water to a total volume of 10 μL per sample. Primers (**Supplementary Table 4**) were purchased from Sigma Aldrich; the reverse Primers were tagged at their 5’ end with fluorescent tags (HEX, FAM or JOE). The first multiplexing reaction contained primers GT023 (HEX), EV037 (HEX) (FAM) in forward and reverse direction. The multiplexing conditions were denaturation at 95°C for 15:30 minutes, annealing 1:30 minutes at 59°C, elongation at 72°C for 11:30 minutes and held at 12°C. The second multiplexing reaction contained primers GT575 (HEX), GATA028 (FAM) in forward and reverse direction, the multiplexing conditions were denaturation at 95°C for 15:30 minutes, annealing 1:30 minutes at 54°C, elongation at 72°C for 11:30 minutes and a final temperature of 12°C. The third multiplexing reaction was set up with primers GATA098 (FAM), GT509 (FAM) and GATA417 (JOE) in forward and reverse direction. The multiplexing conditions were denaturation at 95°C for 15:30 minutes, annealing 1:30 minutes at 54°C, elongation at 72°C for 11:30 minutes and a final temperature of 12°C. The fourth multiplexing reaction contained primers GT310 (HEX) and EV001 (JOE) in forward and reverse direction. The multiplexing conditions were denaturation at 95°C for 15:30 minutes, annealing 1:30 minutes at 60°C, elongation at 72°C for 11:30 minutes and a final temperature of 12°C. After amplification, the multiplexing reactions were diluted with 100 μl water (HPLC grade). Genescan ROX^500^ dye size standard (Thermo Fisher Scientific) and HiDi Formamide (Thermo Fisher Scientific) were mixed 1:100 and used for sequencing in a mix of 1 μl of diluted multiplexing product and 10 μl HiDi-ROX^500^. STR Multiplex fragment analysis was carried out on the 16-capillary electrophoresis system ABI 3300 Genetic Analyser (Applied Biosystems).

#### STR analysis

We first performed MSA analysis [33] using standard parameters, which calculated Weinberg expectation (Fis), Shannon Index (Hs), allele numbers (A) and allele sizes. To detect both weak and strong population structure, we ran simulations with LOCPRIOR and USEPOPINFO, respectively. For both simulations, the more conservative “correlated allele frequencies” -model was used, which assumes a level of non-independence. To ensure that a sufficient number of steps and runs have been performed we used a burn-in period of 1.000 and runs of 100.000 MCMC repeats for both types of simulations, each for 50 iterations for successive K values from 1 - 10. [34]. We used the *a-priori* location to be able to detect even weak population differentiation. In both datasets, we used the web-based STRUCTURE Harvester software [35] to determine the rate of change in the log probability between successive K values, via the *ad-hoc* statistic ∆K from [36]. Figures were rendered using STRUCTURE PLOT V2.0. [37]

### Sequencing of the Mitochondrial Hypervariable Region on the D-loop

The noncoding mtDNA-D-Loop region of 143 individuals was amplified in two overlapping PCR-reactions. We PCR amplified two different lengths fragments for each individual: 1066 bp and 331 bp and sequenced the longer product in forward and the shorter in the reverse direction. The used primers (found in **Supplementary Table 4**) were MT4 (M13F) and MT3 (M13R) for the longer product, and BP15851 (M13F) and MN312 (M13R) for the shorter PCR product from [38]. The 10 μL reaction was carried out with 40 ng genomic DNA and 0.2 μM of each primer and 5 mM Multiplex PCR Kit (Qiagen). Cycling conditions were identical for both directions with 95°C 15:30 min, 53°C 1:30 min, 72°C 13:30 min and hold at 4°C in a Veriti Thermal Cycler (Applied Biosystems). The PCR products were purified with 3 μl Exo/SAP and then cycle-sequenced with the BigDye™ Terminator v3.1 Cycle Sequencing Kit (Thermo Fisher Scientific) according to manufacturer instructions with BP15851 (M13F) for the forward PCR product and MN312 (M13R) for the reverse PCR product, respectively. The mixes were then purified with BigDye X-Terminator™ Purification Kit (Thermo Fisher Scientific) and sequenced by capillary electrophoresis on an ABI 16-capillary 3130xl Analyzer (Applied Biosystems). The sequences were *de-novo* assembled, and consensus sequences were generated with Geneious Software 10.2.3 [39].

### Prdm9 coding minisatellite array PCR and Sequencing

To characterise the minisatellite-coding for the C_2_H_2_-ZnF array in more detail, we designed primers nested between the first two conserved ZnFs, and the reverse primer identical to the long-range amplification primer distal to the coding sequence for the C_2_H_2_-ZnF array. The minisatellite coding for the C_2_H_2_-ZnF array of PRDM9 of 143 *Balaenoptera* individuals was amplified from 20 ng genomic DNA, in a 20 μl PCR reaction, optimised for the amplification of minisatellites. With 0.5 mM primers that were designed for this study using the *Balaenoptera. acutorostrata scammoni (*XM_007172595) reference and included the PRDM9 minisatellite-like C_2_H_2_-ZnF array and additional 100bp flanking regions at 5’ and 3’. Primers Prdm9ZnFA_Bal_R: and Prdm9ZnFA_Bal_F (**Supplementary Table 4**) and 1x AJ-PCR Buffer described in [40] and 0.025 U/μl Taq-Polymerase and 0.0033 U/μl Pfu-Polymerase. Cycling conditions were: initial denaturation at 95°C for 1:30 min followed by 33 cycles including 96°C 15 s, 61°C 20 s and 70 °C 2:00 min and finally 70°C 5 min, and hold at 4°C in a Veriti Thermal Cycler (Applied Biosystems). Agarose gel electrophoresis (1.5% Top Vision Low Melting Point Agarose gel (Thermo Fisher Scientific) with SYBR Safe (Thermo Fisher Scientific) was used to visualise allele sizes as well as zygosity. All bands were excised from the gel (Molecular Imager® Gel Doc™ XR System with Xcita-Blue™ Conversion Screen (Biorad), and PCR products were recovered from the gel fragments with 2 U/100 mg Agarase. If the individuals were homozygous, the extracted DNA was directly Sanger sequenced from in 5’ and 3’ directions for each sample with the BigDye™ Terminator v3.1 Cycle Sequencing Kit (Thermo Fisher Scientific) according to the manufacturer’s protocol with the same primers as for the amplification (Prdm9ZnFA_Bal_R/Prdm9ZnFA_Bal_F). The sequencing-reaction was carried out in the ABI 16-capillary 3130xl Analyzer (Applied Biosystems). Heterozygous samples were subcloned before sequencing.

### Subcloning of heterozygous alleles of identical lengths

Subcloning was performed for a subset of samples when Sanger sequencing revealed heterozygous alleles of the same length, that could not be distinguished by electrophoresis, but revealed heterozygous nucleotides in the chromatogram. Instead of inferring haplotypes, we wanted to experimentally ascertain both alleles. Thus, we cloned the remaining PCR product into the TOPO TA (Invitrogen) vector and transferred the vectors into OneShotTop10 chemically competent cells (Invitrogen). All steps were carried out according to the manufacturer’s manuals. Eight positive clones were picked for each sample, and the DNA was extracted by boiling in HPLC-grade water at 96°C for 10 Minutes. A centrifugation step then removed the cell debris, and the supernatant was directly used for PCR, gel-purification and Sanger-sequencing as described in the section above.

### Sequence Analysis of the PRDM9 C_2_H_2_-ZnF array

Sequences determined different alleles and numbers of repeat units per array, which were *de novo* annotated and the resulting consensus sequences were translated into the corresponding protein variants of each C_2_H_2_-ZnF by Geneious Software 10.2.3 [39]. All PCR products contained the complete minisatellite-like coding sequence for the C_2_H_2_ domain and additional 100 base pairs flanking regions at the 5’ and 3’ ends.

### PRDM9 C_2_H_2_-ZnF array coding sequence dN/dS analysis

The internal C_2_H_2_-ZnFs were obtained as described for the phylogenetic reconstruction. Subsequently, all repeat units were stacked using only non-unique alleles. The resulting codon alignment was used to determine basic population parameters for each population (population theta using segregating sites and nucleotide diversity) by either including or excluding hypervariable amino acid positions (−1, +3, +6) with the R package “pegas” [41]. Episodic diversifying selection among Zinc-finger sites using was determined by a mixed-effects model of evolution (MEME) at https://www.datamonkey.org as described earlier [42].

### Phylogenetic analyses of the PRDM9 hypervariable domain

The gene tree of PRDM9 was inferred either using only the repeats coding for the internal C_2_H_2_-ZnFs or the concatenated KRAB, SSXRD and SET domain. For the internal C_2_H_2_-ZnFs based phylogenetic reconstruction (**Figure 1**), internal C_2_H_2_-ZnFs were extracted, and pairwise distances were calculated as in Vara et al. 2019 [43] with the developmental R package repeatR (https://gitlab.gwdg.de/mpievolbio-it/repeatr). In brief, only species were used with one leading C_2_H_2_-ZnF and at least eight internal C_2_H_2_-ZnFs (see supplementary Figure 1). For each possible repeat combination (r, r’) the hamming distances of the corresponding repeat units r = (r_1_; r_2_; r_3_; …) and r’ = (r’ _1_; r’ _2_; r’ _3_; …) were used to derive the edit distance between r and r’. Before calculating the edit distance, the codons coding for the hypervariable amino acid positions (−1, +3, +6) were removed for each repeat unit and weighting cost of w_mut_ = 1, w_indel_ = 3.5 and w_slippage_ = 1.75 as given in [43]. The resulting pairwise edit distance matrix was used to calculate a neighbour-joining tree with the *bionj* function of the R package ape [44], rooted on the branch leading to the bottlenose dolphin (*Tursiops truncatus)* and visualized in Figtree (http://tree.bio.ed.ac.uk/software/figtree/ v1.4.4).

**Figure 1.**
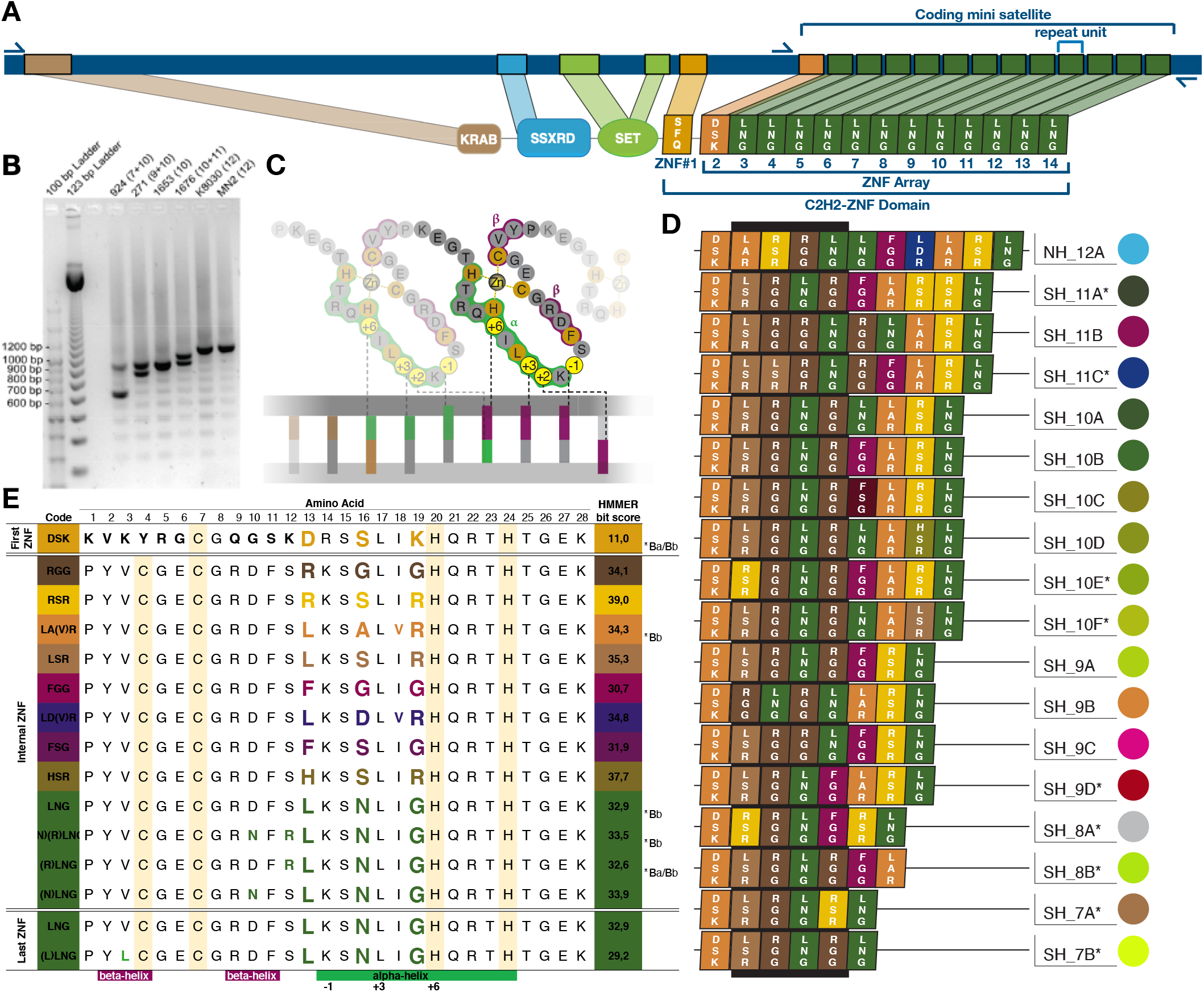
Diversity in the Cys2His2-ZnF domain in minke whales. (A) *Prdm9* gene annotation in *Balaenoptera acutorostrata scammoni* genome reference, with primer site annotations. We predicted the PRDM9 protein (B) Representative PCR products of DNAs from individuals from AN IV (294,271), AN V (1653), NP (K8030) and NA (MN2) showing variation in the number of minisatellite repeats units (number of ZnFs in brackets) (C) Stylized structure of C_2_H_2_-ZnF binding to DNA, with nucleotide-specificity conferred by amino-acids in positions −1, +2, +3, and +6 of alpha-helices. (D) PRDM9 C_2_H_2_-ZnF arrays identified in minke whales, named with broad-scale sampling location, Northern Hemisphere (NH) and Southern Hemisphere (SH) and total number of ZnFs in the C_2_H_2_-ZnFs domain. (E) Types of PRDM9 C_2_H_2_-ZnFs in minke whales. We generate three-letter-codes using the IUPAC nomenclature of amino-acids involved in DNA-binding. All variable amino acids are colored, and asterisks label C_2_H_2_-ZnFs also present in genome references of *Balaenoptera acutorostrata scammoni* (*Ba) or *Balaenoptera bonarensis* (*Bb), see also Supplementary Figure 4.

### Prediction of DNA-binding motifs of different PRDM9 variants

DNA-binding Specificities of the different Cys_2_His_2_ Zinc Finger Protein variants were predict *in-silico* using the SVM polynomial kernel method within “Princeton ZnF” (http://zf.princeton.edu/) [45].

## Results

### The evolutionary context of PRDM9 in *Cetartiodactyla*

For a broad view on the evolution of PRDM9 in *Cetartiodactyla*, we identified PRDM9 orthologs from all publicly available genomic resources (**Supplementary Table 2**). To first establish an evolutionary context based on the protein-coding regions for the proximal conserved domains, we used the *Balaenoptera acutorostrata scammoni* protein as a query and extracted coding sequences of the N-terminal (conserved) domains. Previous studies of PRDM9 in vertebrates found rapid evolution of DNA binding sites only when the KRAB and SSXRD domains were present [46]. Consequently, we used InterProScan to create a curated dataset of *Cetartiodactyla* PRDM9 orthologues which contained the KRAB, SSXRD, and SET domains, including a PRDM9 orthologue in Antarctic minke whale. Our initial analyses revealed that minke whales, and *Cetartiodactyla* in general, possessed all the features characteristic of organisms with PRDM9-directed recombination initiation, including complete KRAB, SSXRD and SET domains. However, organisms with PRDM9 directed recombination, also display a rapidly-evolving PRDM9 C_2_H_2_-ZnF array [46] With the sole exception of Hippopotamus, all *Cetartiodactyla* also contained the C_2_H_2_-ZnF array, with large variation in the number of C_2_H_2_-ZnF between species.

### Characterizing the *Prdm9* gene in minke whales

To characterize the sequence and structure of the *Prdm9* gene, we began by amplifying and sequencing it from a pooled sample, containing DNA from six individuals, five Antarctic minke whales and one common minke whale. The consensus sequence was subjected to *in-silico* prediction that successfully recovered all relevant PRDM9 protein domains with high-confidence: the KRAB domain (E-value: 1.84e^−11^); the SSXRD motif (E-value: 4.46e^−10^); the PR/SET domain (E-value: 9.69e^−05^); and several Zinc-Fingers (**Figure 1**). The complete C_2_H_2_-domain comprises an array of C_2_H_2_-ZnFs (the C_2_H_2_-ZnF-array), as well as a single C_2_H_2_-ZnF (ZnF#1) that is located proximally (E-value: 6.52e^−04^). This PRDM9-ZnF#1 is identical to that of mice [47] as well as rats, elephants, human, chimpanzee, macaque and orangutan [48], and thus appears to be conserved across large evolutionary timescales. The second C_2_H_2_-ZnF (E-value: 1.65e^−11^) appears conserved across *Balaenoptera*. The same second C_2_H_2_-ZnF is seen in our pooled sample, as well as the genome reference of the North Pacific minke whale (*Balaenoptera acutorostrata scammoni)* and Antarctic minke whale (*Balaenoptera bonarensis)* as well as the blue whale (*Balaenoptera musculus).* However, from the third ZnF onwards, sequencing across pooled samples exhibited high variability, which we could not resolve.

### Variation of the PRDM9 coding minisatellite in minke whales

We assessed the variability of the C_2_H_2_-ZnF-array by analyzing the coding minisatellite in a large number of minke whales including Antarctic minke whale (*Balaenoptera bonaerensis* Burmeister, 1867) and two Common minke whale subspecies, Atlantic minke whale and North Pacific minke whale. Amplification and subsequent electrophoresis on agarose gels resolved six different allele sizes of the last exon of the *Prdm9* gene. A single size was observed in all Common minke whales from sampled populations in the North Atlantic as well as North Pacific. In contrast, five different sizes were identified across Antarctic minke whale populations sampled in the Southern Hemisphere. All common minke whales have a size consistent with eleven 84 bp repeats while the most prevalent genotype in the Antarctic minke whales corresponds to nine repeats, with variation between six and ten repeats. Therefore, *Prdm9* shows size variation length of the coding minisatellite, that result from variation in the number of repeat units.

Sequencing of the DNA purified from gel fragments, as well as cloning of apparently homozygous bands, revealed a remarkable diversity of minisatellite alleles. Over 80% of animals possessed repeat-arrays of nine repeats; however, these comprised six different alleles. Because in addition to variation in repeat-number, there is nucleotide variation between 84bp-minisatellite repeats. And finally, not only which repeat-types are present, but also their order varies between individuals.

### Diversity and diversifying selection on C_2_H_2_-zinc fingers of PRDM9

Each 84bp repeat unit translates into 28 amino-acids coding for a single C_2_H_2_-ZnF. To explore diversity between the ZnF, we extracted all 84bp repeat units, which we used to predict C_2_H_2_-ZnFs through an HMMER algorithm [45]. This yielded 26 different versions with HMMER bit scores >17 (**Figure 1** and **Supplementary Figure 4**). Based on amino acid variation within each predicted C_2_H_2_-ZnFs, we identified 14 “common” variants found in multiple individuals as well as twelve “rare” variants found in only a single individual (**Supplementary Figure 4**). The most variable amino acids are 13, 16 and 19, which correspond to positions −1, +3 and +6 of the alpha-helices that are responsible for DNA binding specificity. However, five of the fourteen “common” variants have the same amino sequence LNG at these positions, differing instead at three positions in beta-helices flanking the cysteine residues that bind the zinc-ion (positions 3, 10 and 12). Three of the five LNG variants are also found in the reference sequences of both the Antarctic minke whale and the North Atlantic minke whale. We next used the fourteen common C_2_H_2_-ZnFs to test for signals of selection, finding Evidence that amino acid 16, is under episodic diversifying selection (**Supplementary Figure 5**). This amino acid is located at position +3 of the alpha helix, one of the three positions responsible for DNA-binding specificity.

### Evolutionary turnover of PRDM9 in minke whales

Using all minisatellite-coding sequences for the array, we determined C_2_H_2_-ZNF domain variants that each individual possessed. Minisatellite size homoplasy equates to an identical PRDM9 C_2_H_2_-ZnF array in Common minke whales, but not Antarctic minke whales. The only C_2_H_2_ domain variant found in all samples from the Northern Hemisphere NH12_A consists of eleven C_2_H_2_-ZnF in the array and the C_2_H_2_-ZnF, that is located proximally and appears conserved across *Balaenoptera*. In contrast, samples from the Southern Hemisphere displayed seventeen different C_2_H_2_-Domains of PRDM9 (**Figure 1**).

We want to understand the evolutionary relationships of the different alleles that we observed in minke whale species. However, phylogenetic analysis of minisatellite repeat-structures is challenging due to their rapid rate of evolution. Minisatellites, including that of *Prdm9*, evolve mainly by unequal crossing-over and gene conversion in meiosis [49]. Stepwise mutation models that are commonly assumed are based on microsatellites, and thus give only a poor fit to minisatellite evolution [50]. So, instead of comparing full-length-*Prdm9* allelic variants, we separated each array into 84bp repeat types. We then calculated pairwise Hamming (i.e. minimum edit) distances between all repeats, after removing the nucleotides coding for the hypervariable amino acid positions (−1, +3, +6), and applying specific weighting costs as given in [43]. In addition to the repeats present in our minke whale samples, we also included repeats coding for the internal C_2_H_2_-ZnFs of all *Cetartiodactyla* species into our phylogenetic analyses (see **Supplementary Figure 1**) Here, we included only those species that possessed the complete N-terminal domain architecture, the proximal C_2_H_2_-ZnF and at least eight successive C_2_H_2_-ZnF in the C-terminal DNA-binding domain. We restricted our phylogenetic analyses on the internal minisatellite repeats since our analyses revealed that the DNA-binding positions of the most proximal ZnF in the array (ZnF#2) are not only conserved in minke whales but also generally within *Cetacea* (**Supplementary Figure 2**). In fact, we observed only a single DNA-binding amino-acid change between C_2_H_2_-ZnF #2 of all *Cetacea* compared to all *Ruminantia* (**Supplementary Figure 3**).

The PRDM9 phylogenetic analyses in **Figure 2** show that all species cluster into distinct phylogenetic groups. Reference alleles cluster with their own subspecies, even including the unusual reference allele of *Balaenoptera acutorostrata scammoni*, which nevertheless clusters with our common minke whale samples. Similarly, the genome reference allele of *Balaenoptera bonarensis* (Antarctic minke whale) clusters within the group of SH_8A and SH-8B alleles found in our Antarctic minke whale samples, which appear to be more ancestral than other minke whale alleles. The phylogenetic reconstruction also suggests that common minke whale alleles have appeared more recently and evolved mainly by an increase in repeat-copy after splitting from the Antarctic minke whale. Our phylogenetic analysis thus fits well with an evolutionary history where common minke whales split from Antarctic minke whales approximately 5 million years ago.

**Figure 2.**
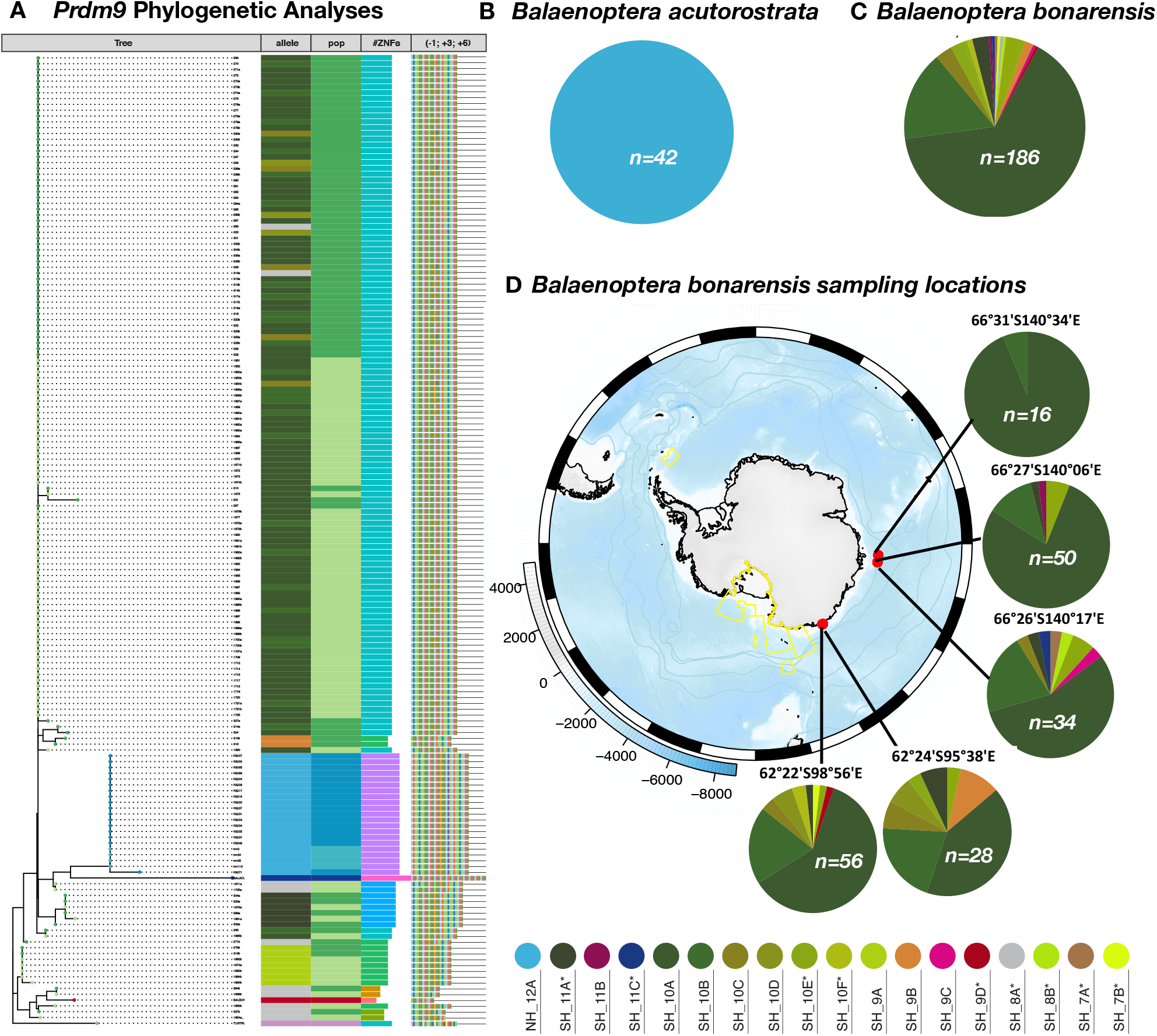
*Prdm9* Phylogenetic analyses and allele frequencies at different geographical scales. (A) PRDM9 phylogenetic analyses across all geographical regions, including several outgroups, from top to bottom: **−1, +3, +6**: PRDM9 variants as in Figure 1. **# ZnFs:** number of ZnFs **pop:** assigned population of individuals (dark blue): North Pacific, (light blue) North Atlantic, (light green) Antarctic Area IV, (dark green) Antarctic Area V, as well as Genome Reference alleles: (blue*) Balaenoptera acutorostrata scammoni*, (red) *Balaenoptera bonarensis* and outgroups (pink) *Tursiops Truncatus* **allele:** PRDM9 coding minisatellite allele colored as the translated variant from Figure 1. **Tree:** Nucleotide Phylogeny of PRDM9 alleles. (B) allelic diversity in Common minke whales (C) allelic diversity in Antarctic minke whales (D) fine-scale diversity by Sampling locations of Antarctic minke whales on a map of the Southern Ocean (yellow delineations show protected areas).

### Diversity of PRDM9 at different geographical scales

To explore the allelic diversity of *Prdm9* at different geographical scales, we partitioned the data by sampling location wherever accurate catch-locations were available (**Figure 2**). Allelic diversity varied between sampling locations, even within the Antarctic sampling sites. The most common alleles are SH_10A, with SH_10B, and these occur at frequencies of 10-20% in all sampling locations. However, all other alleles differ between different sampling populations, each of which carries at least one allele found exclusively in a single population. Sample size does not predict allele number, with the highest allele number (ten) being associated with an average-sized sample (n=34).

### Population structure of minke whales

We are interested in the possible evolutionary consequences of PRDM9 for minke whale speciation in the face of what may be recent secondary mixing, and this requires a context of current levels of population isolation. We thus explored variability at the 84bp *Prdm9* minisatellite repeat across the four sample regions for population genetic analyses (**Table 1**). Taking each repeat type as an individual allele, we computed the mutation rate per base pair per generation (population θ), as well as the average pairwise nucleotide diversity. The latter was calculated in two ways: (i) using the entire minisatellite-like repeat sequence; (ii) after removing nucleotides coding for the hypervariable sites that translate into the DNA binding positions of individual C_2_H_2_-ZnF, which are known to be under positive selection. We observed the highest population θ in Antarctic Area IV, compared to all other sampling locations, as seen in **Table 1**. Across the microsatellite-repeat types, the Average Pairwise Nucleotide Diversity (APND, including nucleotides coding for the hypervariable sites) is similar across all sampled populations and between common minke whale and Antarctic minke whales. After excluding the nucleotides coding for hypervariable sites (−1, +3, +6), the APND is decreased roughly 2.5-fold, across minke whales.

**Table 1.**
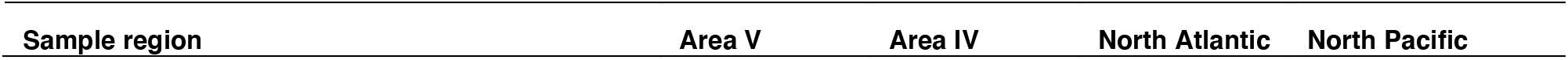
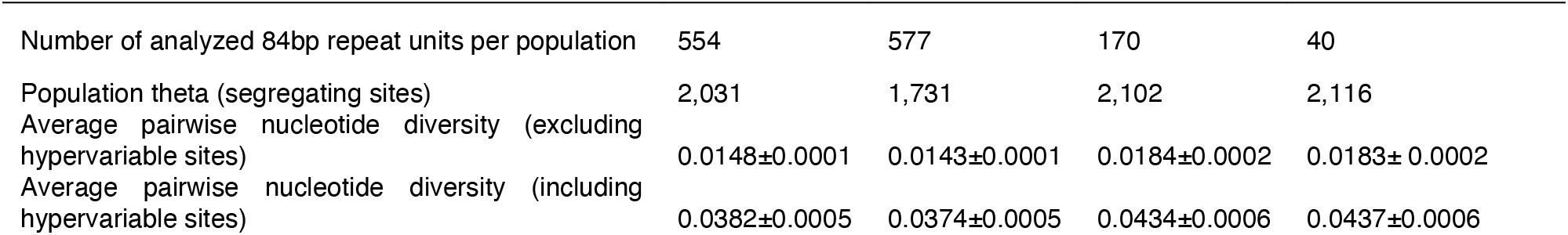
**Nucleotide diversity [68] of the minisatellite coding for the PRDM9-C2H2-ZnF array**, analyzed per sample region. Average pairwise nucleotide diversity was analyzed for the all sites, and also excluding the nucleotides coding for amino-acids at hypervariable sites (−1, +3, +6) in the alpha-helix of C_2_H_2_-ZnFs.

| Sample region | Area V | Area IV | North Atlantic | North Pacific |
| --- | --- | --- | --- | --- |
| Number of analyzed 84bp repeat units per population | 554 | 577 | 170 | 40 |
| Population theta (segregating sites) | 2,031 | 1,731 | 2,102 | 2,116 |
| Average pairwise nucleotide diversity (excluding hypervariable sites) | 0.0148±0.0001 | 0.0143±0.0001 | 0.0184±0.0002 | 0.0183± 0.0002 |
| Average pairwise nucleotide diversity (including hypervariable sites) | 0.0382±0.0005 | 0.0374±0.0005 | 0.0434±0.0006 | 0.0437±0.0006 |

We used two approaches to quantify genetic differentiation of PRDM9 between the different minke whale populations: the per-site distance for multiple alleles [51] and Jost’s D (D_J_), the fraction of allelic variation among populations [52]. Both measures indicate weak population differentiation, the highest differentiation being between the North Atlantic and North Pacific (G_ST_ = −0.0039, D_J_ = 0.0077). The second highest population differentiation was seen between the North Pacific and both Antarctic regions, G_ST_ = −0.0035, D_J_ = 0.0070, with slightly lower differentiation between Atlantic and Antarctic, G_ST_ = −0.0011, D_J_ = 0.0022. The lowest population differentiation was between Antarctic Areas IV and V (ANV vs ANIV, G_ST_ = −0.0004, D_J_ = 0.0009).

To explore population structure differentiation further, we also analysed polymorphic microsatellites and the hypervariable region of the mitochondrial D-Loop (mtDNA HVR) (**Figure 3**). Despite the low *Prdm9* diversity in the Northern Hemisphere, genetic structuring was revealed by the mtDNA HVR, with a clear distinction between the North Atlantic and North Pacific whales. We also examined nine unlinked autosomal microsatellite loci, chosen for their high information content in minke whales [17, 29] and sex-chromosome markers ZFX/ZFY, from [38]. Microsatellite summary statistics reveal high heterozygosity of effectively zero F_IS_ (Supplementary Table 1), so we continued with a Bayesian analysis of population structure. We used *STRUCTURE* with a recessive allele model because any null alleles were due to polymorphisms, rather than missing data [53] and the “admixture” model. However, given that minke whales occur in geographically isolated groups, are separated by hemispheres, and have asynchronous breeding seasons, we nevertheless assumed a low probability for incomplete lineage sorting, which could otherwise confound structure inference, particularly for weak population differentiation. The most likely number of clusters returned by ΔK is *k* = 2, which distinguishes the two hemispheres. Increasing to *k = 3* and *k = 4*, further separates the common minke whale subspecies (as shown in **Figure 3**), a result that holds even without *a priori* location data (NOLOCS) where only strong population structure should be detected (**Supplementary Figure 6**).

**Figure 3.**
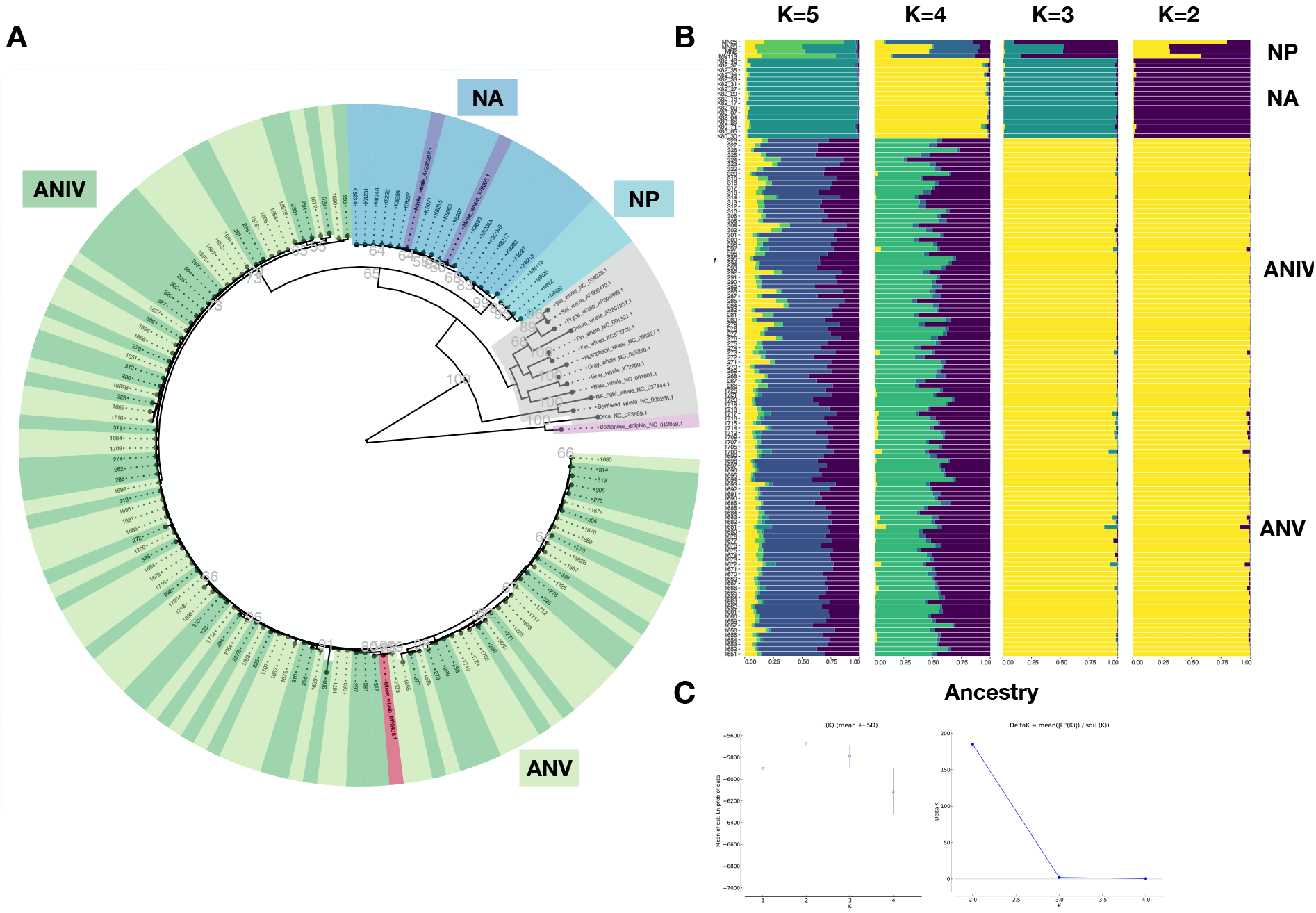
**Population structure of minke whales** (NP) North Pacific, (NA) North Atlantic (ANIV) Antarctic Area IV (ANV) Antarctic Area V (A) Phylogeny of minke whale mitochondrial D-loop region, containing the hypervariable segments (HVS). (B) Population STRUCTURE analyses of minisatellites with *a priori* location information. (C) Log probability of data L(K) as a function of K (mean ± SD) (D) Magnitude of ΔK as a function of K (mean ± SD over 50 replicates).

### Putative minke whale Recombination initiation motifs

Different C_2_H_2_-ZnF domains target different recombination hotspots [3, 4, 54, 55]. Target hotspots in humans [4, 48] and mice [55] can be predicted from C_2_H_2_ ZnF sequences using the SVM polynomial kernel model for *de-novo* binding prediction [45]. To understand the situation in minke whales, we identified a number of “core motifs”, based on ZnFs #3-#6, that appear particularly important for DNA binding [56, 57] (**Figure 4**). As must be true, the single PRDM9 variant found in common minke whales results in a single “core motif”. However, a large majority of Antarctic minke whales also share a single “core motif”, despite a much larger diversity of PRDM9 (**Figure 4**). Core motifs show no overlap between common minke whales and Antarctic minke whales, a trend that is particularly evident at C_2_H_2_-ZnF#3 where it creates distinctive DNA binding motifs. Thus, it appears as if core motifs are shared between the majority of minke whale PRDM9 variants within a given species, but not between species inhabiting different Hemispheres.

**Figure 4.**
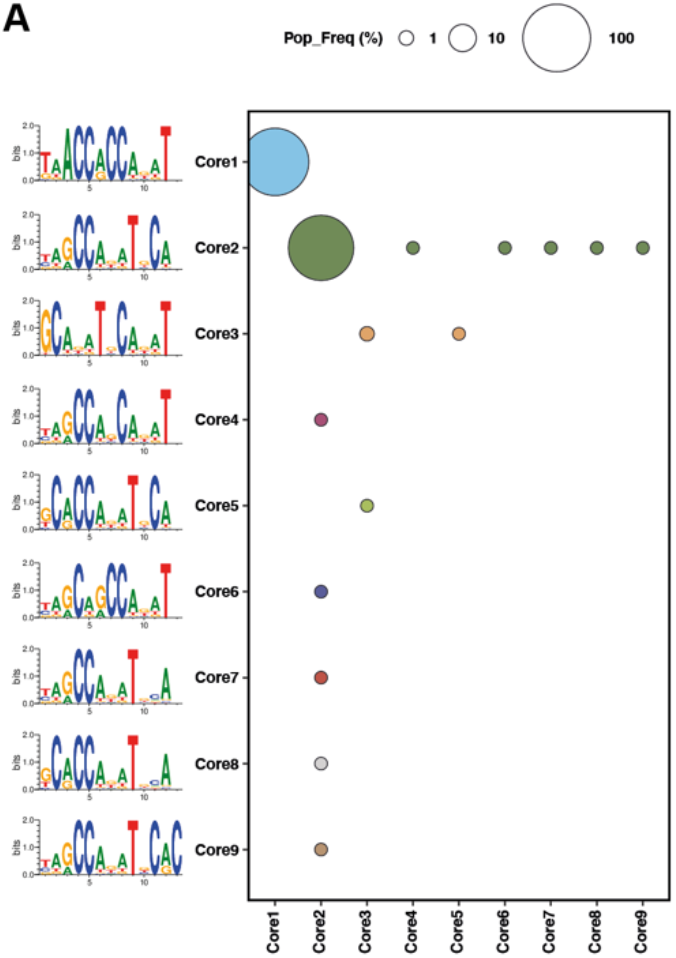
**PRDM9 Array binding predictions** and population frequency based on ZnFs#3-#6 “core motif” combinations across all individuals

## Discussion

### PRDM9 shows high diversity, especially in Antarctic minke whales

We characterized the PRDM9 C_2_H_2_-ZnF domain diversity of 134 individuals from four natural populations of minke whales and discovered a total of eighteen variants. The minisatellite encoding the DNA-binding C_2_H_2_-ZnF array apparently evolves rapidly in minke whales, with at least one of the DNA-recognizing position likely under positive selection. This high diversity is similar to that observed at the *Prdm9* gene in other vertebrate species, such as humans, mice, bovines and primates [4–6, 12, 58].

However, PRDM9 diversity is not equally abundant in both hemispheres. In contrast to the extensive PRDM9 diversity we observe in Antarctic minke whales (*Balaenoptera bonarensis)*, only a single PRDM9 variant is found in Common minke whales. This variant is shared between two subspecies, the North Atlantic minke whale *(B. a. acutorostrata)* and North Pacific minke whales *(B. a. scammoni).* In most mammals, extensive PRDM9 variation typically is seen between, and within, a given subspecies. This holds true also for Antarctic minke whales, thus the observation of an identical PRDM9 variant shared between two subspecies of common minke whales is unexpected. The PRDM9 protein level diversity observed within each Hemisphere contrasted with the levels of population differentiation seen in microsatellite as well as mitochondrial haplotypes. However, diversity between repeat types of the *Prdm9* minisatellite can reveal even weak population structure and is consistent with data on mitochondrial haplotypes and microsatellites. This contrasting pattern between protein-level conservation and nucleotide differentiation is fascinating and points to functional constraints operating on different levels of PRDM9 evolution.

### Population genetic analyses of minke whales and potential speciation

Taxonomists have separated the common minke whale into two subspecies, the North Atlantic minke whale *(B. a. acutorostrata)* and the North Pacific minke whale (*B. a. scammoni*), which diverged approximately 1.5 million years ago. variability between 84bp *Prdm9* minisatellite repeat units, as well as microsatellite markers and mtDNA all reveal weak population structure and differentiation between minke whales in North Atlantic and North Pacific. Our population genetic data support this classification and suggest that these populations may be in the process of speciating. Yet fascinatingly, all individuals of both Northern Hemisphere populations had identical C_2_H_2_ domains, which suggests that these subspecies should still be able to interbreed if given a chance. As geographical isolation of the two subspecies is currently still upheld by the permanent polar ice, allopatric speciation may be promoted. However, due to global warming, the two subspecies might come into secondary contact again in the future. The removal of the current geographical barrier could therefore disrupt speciation, especially in the light of identical C_2_H_2_ domains encoded by the mammalian hybrid sterility gene *Prdm9.*

### Hybrids between Antarctic minke whale and Common minke whale

Analysis of the hypervariable region (HVR) on the mitochondrial D-Loop suggests that Antarctic minke whale and Common minke whale evolved from a common ancestor in the Southern Hemisphere during a period of global warming approximately 5 million years ago [59]. Common minke whales and Antarctic minke whales are now separated by both geography and seasonality. However, while their winter habitats and breeding grounds remain unknown, it is assumed that *B. a. acutorostrata* migrates south between November and March to give birth in warmer waters and occasional individuals are seen as far south as the Gulf of Mexico [60]. Interbreeding between the populations of the Northern and Southern Hemisphere appears unlikely at the time our samples were collected (the 1980s) because our study confirms previous findings [28] that common minke whales and Antarctic minke whales exhibit extensive genetic differences in their mtDNA and microsatellites. However, recent reports have uncovered rare hybridization events between Common minke whale and Antarctic minke whales [16], which shows that interbreeding is unlikely but possible. The question remains whether known reproductive isolation mechanism exists in hybrids between the two minke whale species.

The *Prdm9* gene remains the only hybrid sterility gene known in vertebrates. Large evolutionary turnover of the coding minisatellite is seen in all mammals characterised to date- including minke whales (*this study*). One consequence is genetic variation in the DNA-binding C_2_H_2_-ZnF-domain that in turn introduces variation in species’ recombination landscapes, that differ even within populations of the same species [4, 61]. Based on computed PRMD9 C_2_H_2_-ZnF binding motifs found in Common minke whale and Antarctic minke whale, we would speculate that the recombination initiation landscapes of the two variants would not overlap. The associated variation in recombination landscapes is implicated in generating reproductive isolation in mammals, which has been well characterized in mice. Here, historical erosion of genomic binding sites of PRDM9 ZnF domains has been shown to lead to the introduction of asymmetric sets of DSBs in evolutionary divergent homologous genomic sequences [61, 62]. This asymmetry is likely caused because PRDM9 initiation can erode its own binding via biased gene conversion over long evolutionary timescales.

As a result, in hybrid genomes, the variant of one species preferentially binds the ancestral, binding sites on the homologue of the other species, that have not been eroded, and vice versa. The resulting asymmetry of recombination initiation sites is believed to be responsible for the inefficient DSB repair, defective pairing, asynapsis and segregation of the chromosomes that are observed in intersubspecific mouse hybrids [63–65]. The low PRDM9 diversity in the Northern Hemisphere would predict extensive historical erosion of PRDM9 binding sites in the genomes of Common Minke Whales. A degree of erosion may also be true across the Southern Hemisphere, where most animals share identical core motifs. Thus, the predicted high level of PRDM9 binding site erosion, coupled with non-overlapping minke whale PRDM9 binding motifs would predict asymmetric PRDM9 recombination imitation in minke whale hybrids (**Supplementary Figure 7**), which is implicated in hybrid male sterility.

Despite this potential incompatibility, two viable female minke whale hybrids have been observed, one of which showed signs of successful backcrossing [15, 16]. This observation alone is not sufficient to clarify whether postzygotic isolation mechanisms do or do not generally operate. After all, Haldane’s rule predicts that the sterile sex will be the heterogametic males, rather than the females on which observations were made, an expectation that holds in PRDM9-mediated hybrid sterility in mice [20, 66, 67]. Therefore, it still remains to be understood whether sporadically occurring hybridization events will generate infertile males or, instead, signify that start of a breakdown of genetic isolation. Further studies on minke whale hybrids are necessary to elucidate this matter.

## Conclusion

We uncovered that Minke whales, and related species, possess coding sequences for a complete set of KRAB, SSXRD and SET domains, a necessary feature of organisms with PRDM9-regulated recombination [46]. Sequencing revealed evolutionary turnover of the PRDM9 coding minisatellite. Our phylogenetic analysis revealed a good association of groups with their geographical origin, a high level of size homoplasy in common minke whales and a lower level of size homoplasy in Antarctic minke whales. In Antarctic minke whales (*Balaenoptera bonarensis*), we observe extensive diversity within the C_2_H_2_-ZNF domain of PRDM9, even at fine geographical scale. In direct opposition, we observe no genetic diversity within the C_2_H_2_-ZNF in common minke whales. In each Hemisphere, the PRDM9 protein diversity is contrasting strongly with the underlying population structure we observe between and within subspecies. However, on the nucleotide level, variation between *Prdm9* minisatellite repeat types mirrored the variation in mitochondrial DNA and microsatellites in respective animals. This apparent contradiction points to maintenance of conserved protein sequence, even across minke whale subspecies boundaries.

## Acknowledgements

We thank the Max Planck Society for funding this study.

We thank Nicole Thomsen and Olga Eitel for technical support. We would also like to thank Kevin Glover and Bjørghild B. Seliussen for sharing primer sequences and genotyping conditions for microsatellites markers and the hypervariable segments of the mtDNA D-loop.

## Availability of data and materials

Most data generated or analyzed during this study are included in this published article and its supplementary information files, Genetic data generated and analyzed during the current study are available in the Zenodo repository, https://doi.org/10.5281/zenodo.4309437, and R Scripts are available from https://gitlab.gwdg.de/mpievolbio-it/repeatr.

## Author Contributions

ED designed the experiments, analyzed data and wrote parts of the manuscript. KKU designed experiments and analyzed the data, WA provided the samples and wrote parts of the manuscript and L.O.-H. conceptualized the study, designed the experiments, analyzed data and wrote the manuscript. All authors read and approved the final version of the manuscript.

## Supplementary Information

**Supplementary Figure 1:**
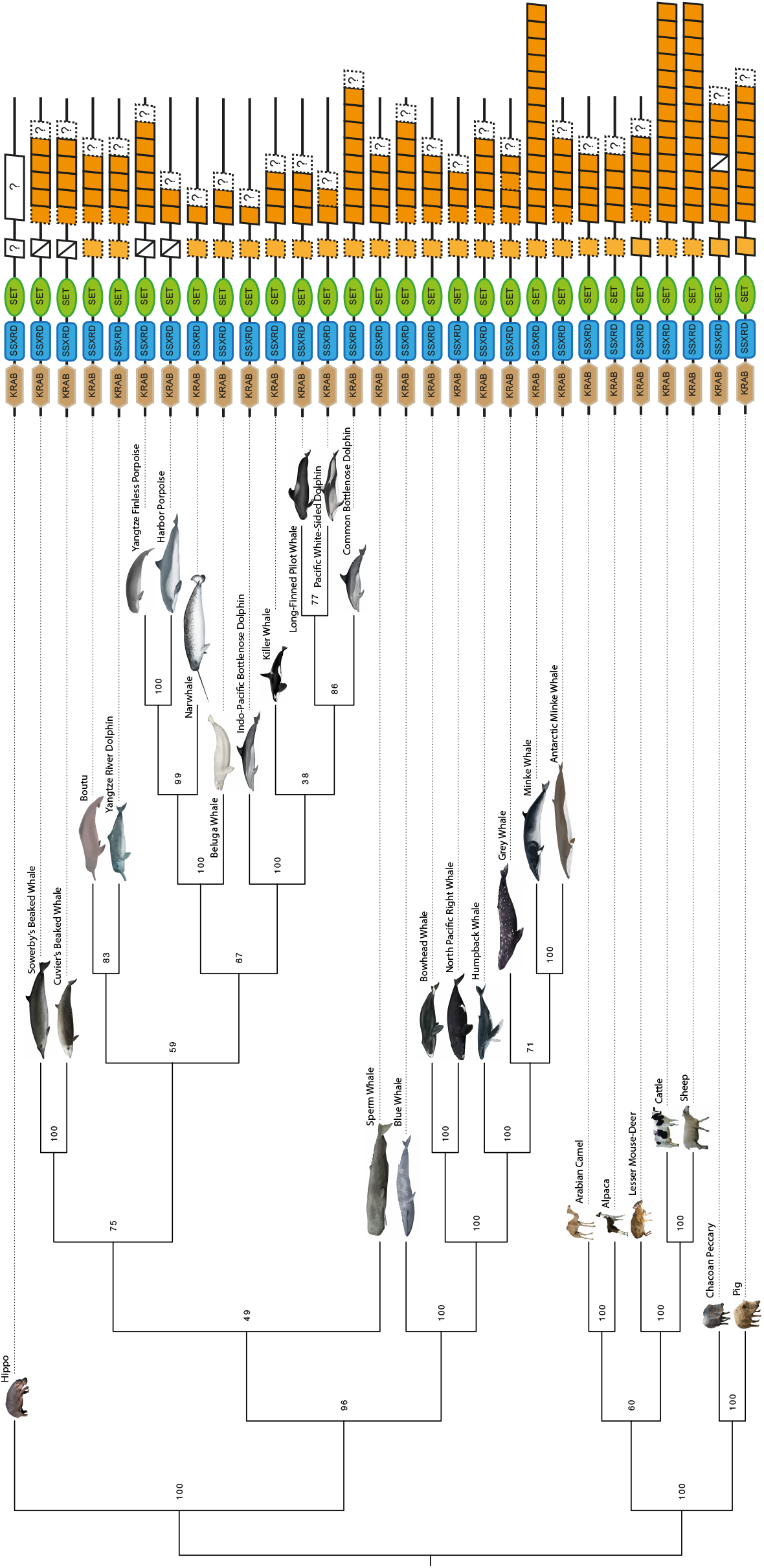
**PRDM9 Phylogenetic analysis across *Cetartiodactyla***, using N-terminal KRAB, SSXRD and SET domains, with data from public genomic databases collected in (**Supplementary Table 2**)

**Supplementary Figure 2:**
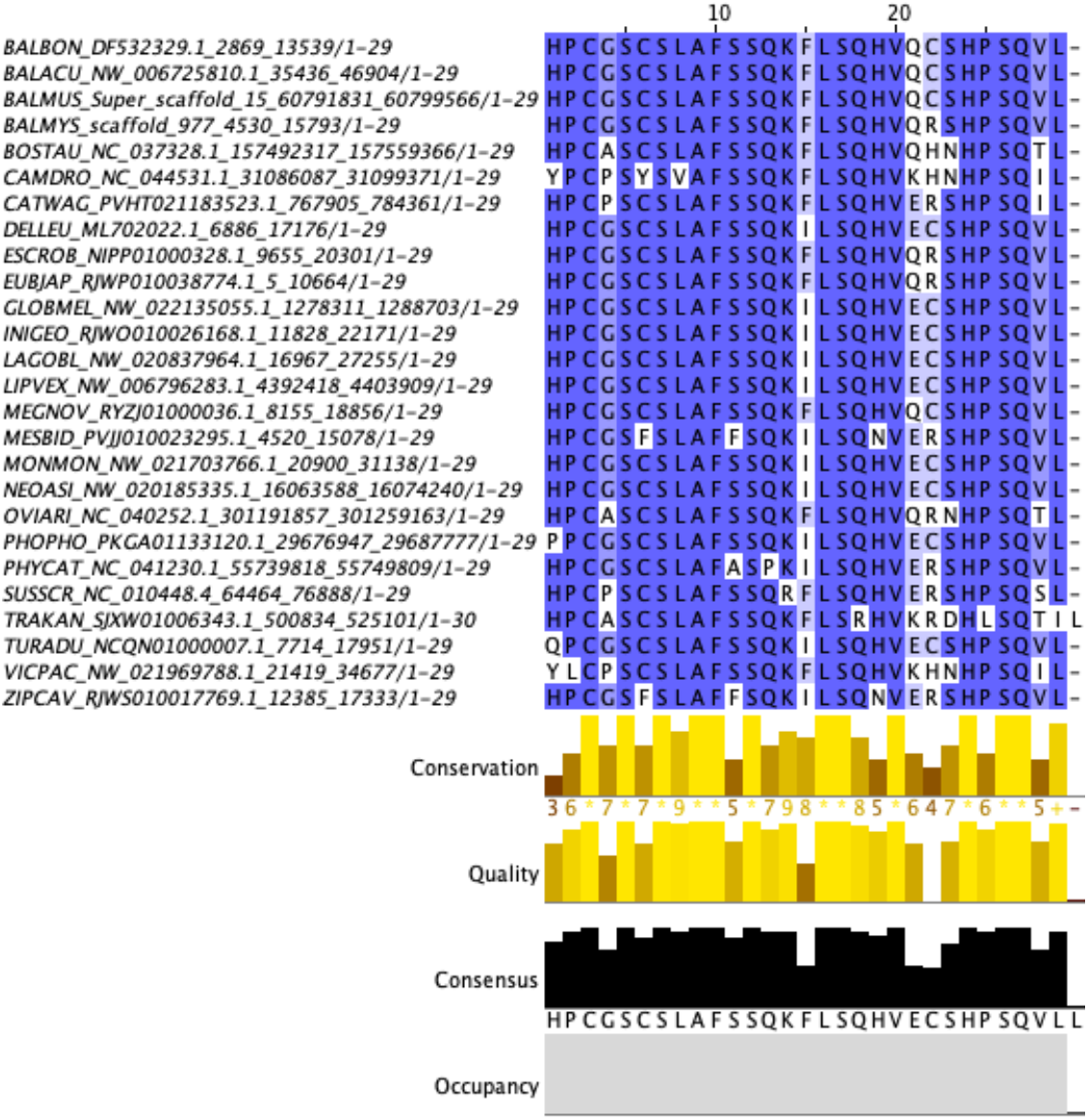
**Alignment of proximal ZnFs of *Cetartiodactyla***, in which this ZnF (outside the ZnF array) was present in the public genomic databases (Supplementary Table **2**). The proximal ZnF is fully conserved across *Balaenoptera.* A single amino-acid change at position 22 in bowhead whale (*Balaena mysticetus)* is observed, compared to *Balaenoptera*, with the DNA contacting residues remaining conserved.

**Supplementary Figure 3:**
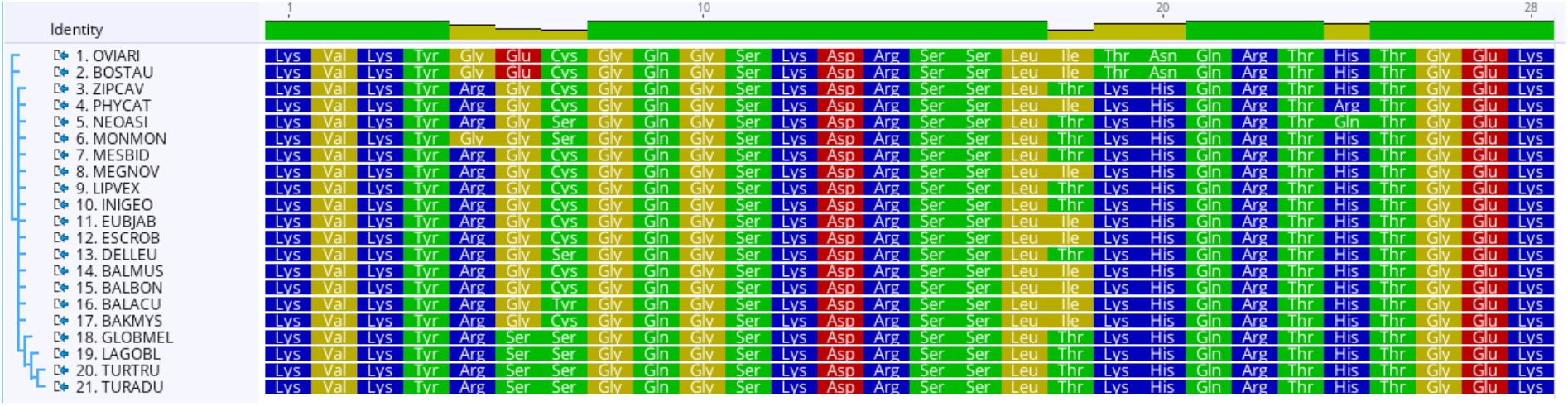
Alignment of second ZnFs of all *Cetartiodactyla*, where this ZnF was present in public genomic databases (Supplementary Table **2**). Across *Cetacea*, ZnF#2 are conserved at the amino-acids 13, 16 and 19, which code for the DNA-binding positions of the alpha-helix of each ZnF. Amino-acids 19 is changed between *Cetacea* and *Ruminantia*, and additional changes are observed at amino-acids 5, 6, 18, and 24.

**Supplementary Figure 4:**
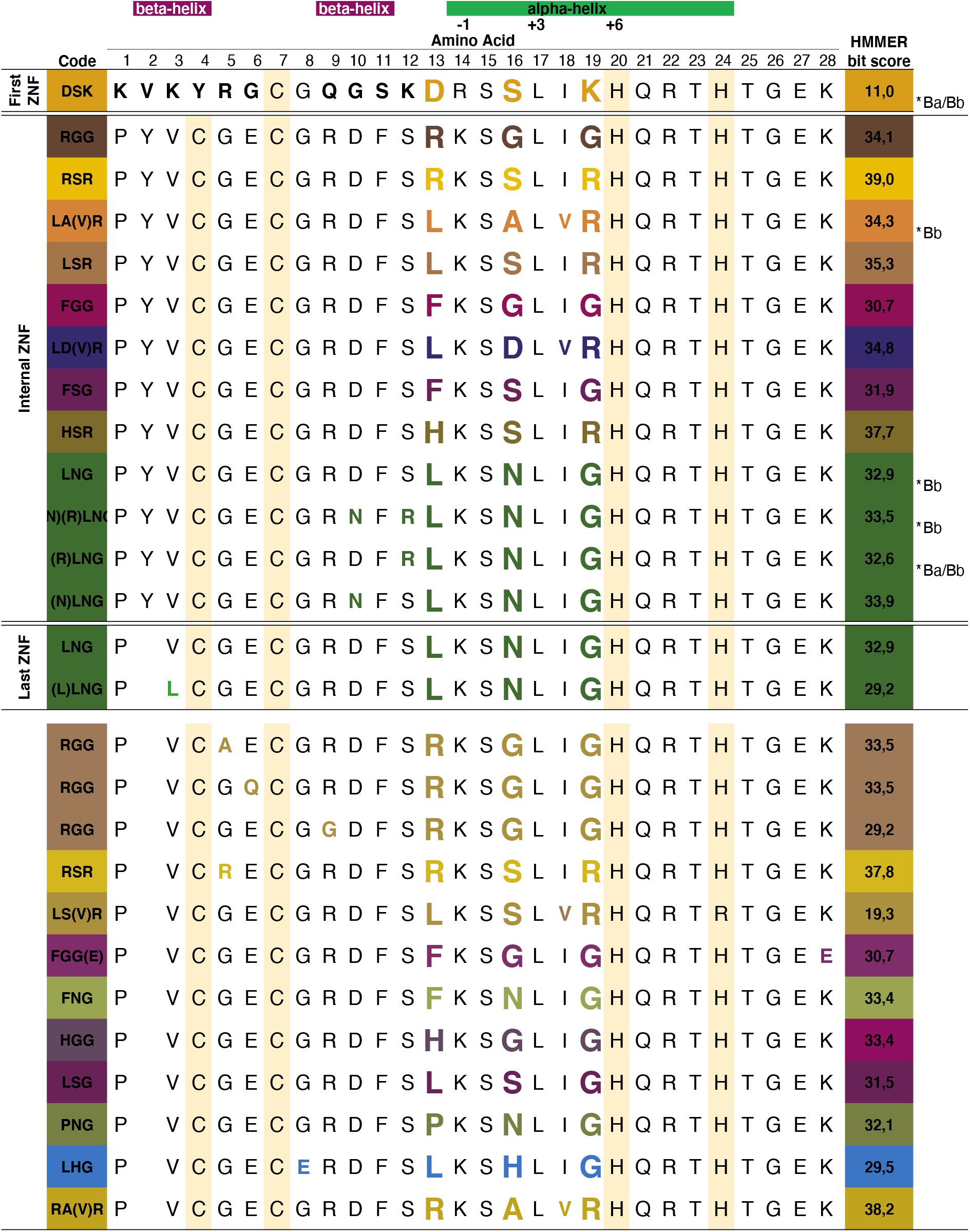
All PRDM9 C_2_H_2_-ZnFs types in minke whales. We generate three-letter-codes for each C_2_H_2_-ZnF using the IUPAC nomenclature of amino-acids involved in DNA-binding. All variable amino acids are colored, and asterisks label C_2_H_2_-ZnFs also present in genome references of *Balaenoptera acutorostrata scammoni* (*Ba) or *Balaenoptera bonarensis* (*Bb).

**Supplementary Figure 5:**
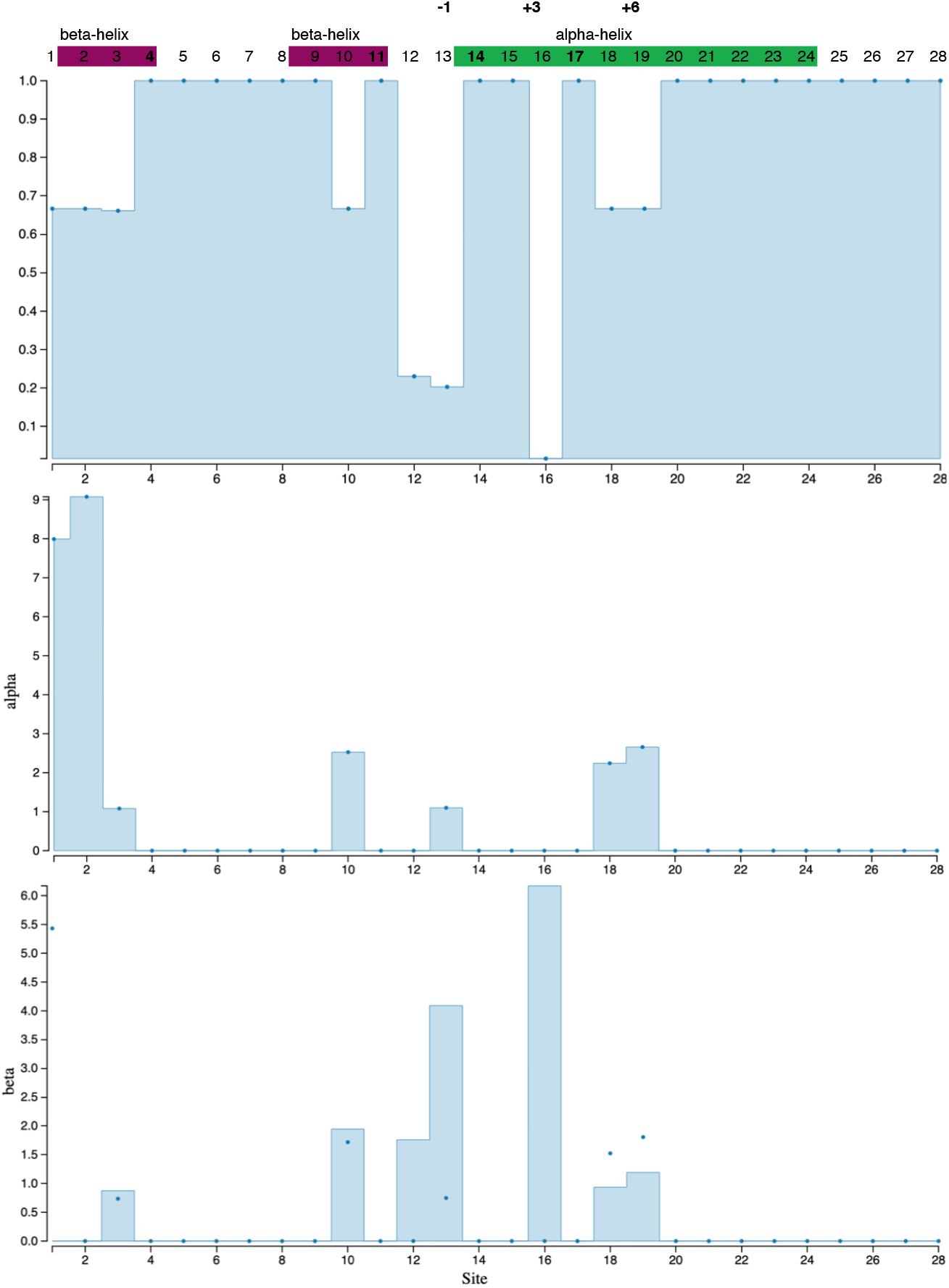
Signals of selection on 84bp nucleotide repeats coding for each ZnF identified in this study. “Site” shows zinc-fingers amino-acid positions, from top to bottom: a site is classified based on whether α>β (negative selection) or α<β (positive selection). (A) p-value is significant when <0,05 (B) fel.α instantaneous synonymous site rate (C) fel.β instantaneous non-synonymous site rate.

**Supplementary Table 1:** Descriptive statistics of microsatellites used in this study analyzed using (Dieringer and Schlötterer 2003). The richness of allelic variants (number of observed alleles) (**A**) Expected Heterozygosity (**H_E_**) observed Heterozygosity (**H_O_**), excess or deficiency from the average number of heterozygotes (**F_IS_**) and (**H_S_**), the probability that two random alleles from the same, randomly chosen subpopulation are different by state.

| Locus | Origin | A | Allele sizes | H<sub>E</sub> | H<sub>O</sub> | F<sub>IS</sub> | H<sub>S</sub> |
| --- | --- | --- | --- | --- | --- | --- | --- |
| GATA028 | AN IV | 20 | 148-272 | 0,93 | 0,74 | 0,21 | 2,73 |
|  | AN V | 20 | 148-274 | 0,92 | 0,88 | 0,05 | 2,67 |
|  | NA | - | - | - | - | - | - |
|  | NP | - | - | - | - | - | - |
| GT575 | AN IV | 26 | 190-262 | 0,95 | 0,95 | 0 | 3,00 |
|  | AN V | 29 | 185-262 | 0,95 | 0,97 | -0,02 | 3,06 |
|  | NA | 8 | 212-240 | 0,85 | 0,89 | -0,06 | 1,89 |
|  | NP | 5 | 206-216 | 0,89 | 1,00 | -0,22 | 1,56 |
| GT310 | AN IV | 18 | 81-135 | 0,88 | 0,78 | 0,11 | 2,41 |
|  | AN V | 18 | 81-137 | 0,90 | 0,82 | 0,09 | 2,46 |
|  | NA | 5 | 113-127 | 0,59 | 0,33 | 0,43 | 1,01 |
|  | NP | 6 | 107-121 | 0,93 | 0,75 | 0,14 | 1,73 |
| EV001 | AN IV | 13 | 103-147 | 0,83 | 0,95 | -0,14 | 1,97 |
|  | AN V | 14 | 105-141 | 0,82 | 0,78 | 0,04 | 1,97 |
|  | NA | 5 | 139-153 | 0,72 | 0,56 | 0,22 | 1,35 |
|  | NP | 4 | 131-151 | 0,86 | 0 | 1,00 | 1,39 |
| GATA098 | AN IV | 9 | 76-108 | 0,83 | 0,89 | -0,08 | 1,85 |
|  | AN V | 10 | 70-108 | 0,84 | 0,97 | -0,16 | 1,94 |
|  | NA | 5 | 80-96 | 0,73 | 0,72 | -0,01 | 1,35 |
|  | NP | 3 | 84-92 | 0,68 | 1,00 | -0,65 | 0,97 |
| GT509 | AN IV | 19 | 179-219 | 0,92 | 0,89 | 0,03 | 2,66 |
|  | AN V | 20 | 179-217 | 0,93 | 0,92 | 0,00 | 2,70 |
|  | NA | 10 | 191-213 | 0,85 | 1,00 | -0,19 | 1,97 |
|  | NP | 5 | 195-211 | 0,89 | 1,00 | -0,22 | 1,56 |
| GATA417 | AN IV | 17 | 188-264 | 0,92 | 0,93 | -0,02 | 2,58 |
|  | AN V | 19 | 184-260 | 0,92 | 0,92 | -0,01 | 2,60 |
|  | NA | 8 | 212-240 | 0,86 | 0,94 | -0,11 | 1,91 |
|  | NP | 5 | 204-242 | 0,89 | 0,75 | 0,11 | 1,56 |
| GT023 | AN IV | 14 | 90-118 | 0,87 | 0,81 | 0,07 | 2,24 |
|  | AN V | 15 | 86-118 | 0,88 | 0,95 | -0,08 | 2,36 |
|  | NA | 8 | 92-108 | 0,77 | 0,78 | -0,02 | 1,64 |
|  | NP | 5 | 94-110 | 0,86 | 1,00 | -0,27 | 1,49 |
| EV037 | AN IV | 18 | 184-222 | 0,91 | 0,49 | 0,46 | 2,53 |
|  | AN V | 16 | 184-222 | 0,93 | 0,52 | 0,44 | 2,59 |
|  | NA | 7 | 192-208 | 0,72 | 0,72 | -0,02 | 1,52 |
|  | NP | 5 | 176-196 | 0,86 | 1,00 | -0,27 | 1,49 |
| Origin |  | A |  | H<sub>E</sub> | H<sub>O</sub> | F<sub>IS</sub> | H<sub>S</sub> |
| Average | AN IV | 17,1 |  | 0,89 | 0,82 | 0,07 | 2,44 |
|  | AN V | 17,9 |  | 0,90 | 0,86 | 0,04 | 2,48 |
|  | NA | 7 |  | 0,76 | 0,74 | 0,03 | 1,58 |
|  | NP | 4,75 |  | 0,86 | 0,81 | -0,05 | 1,47 |

**Supplementary Table 2:**
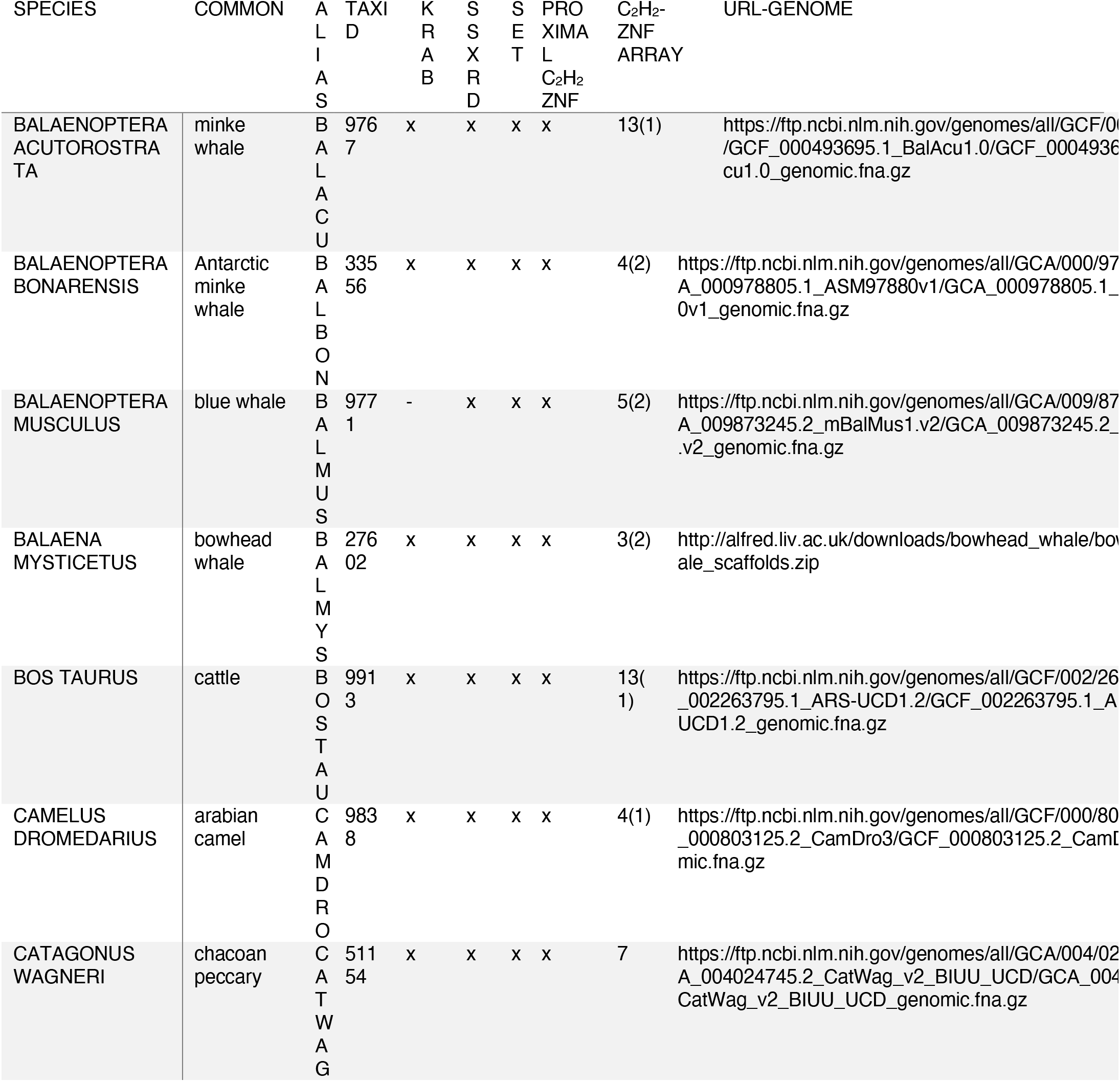
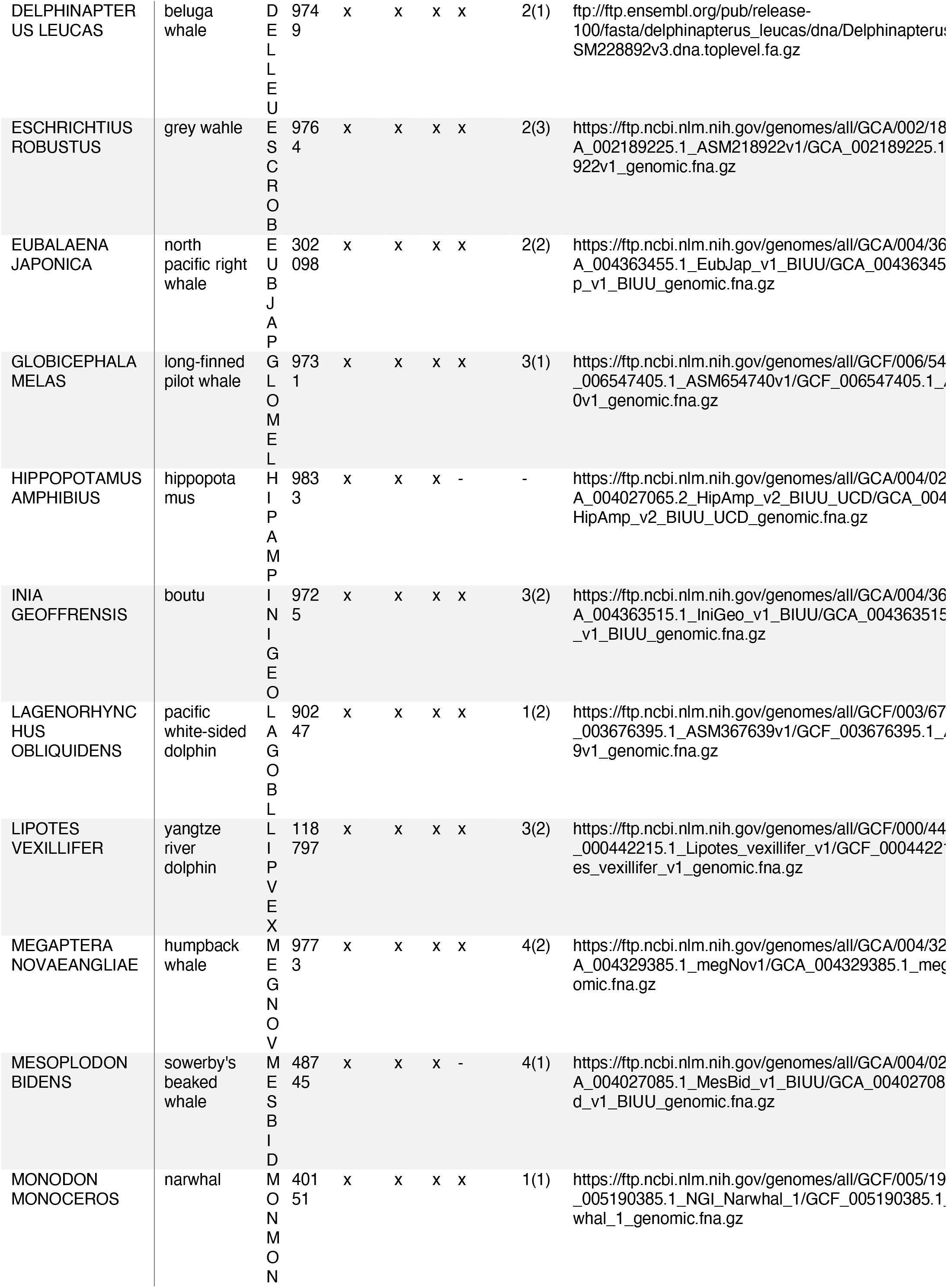
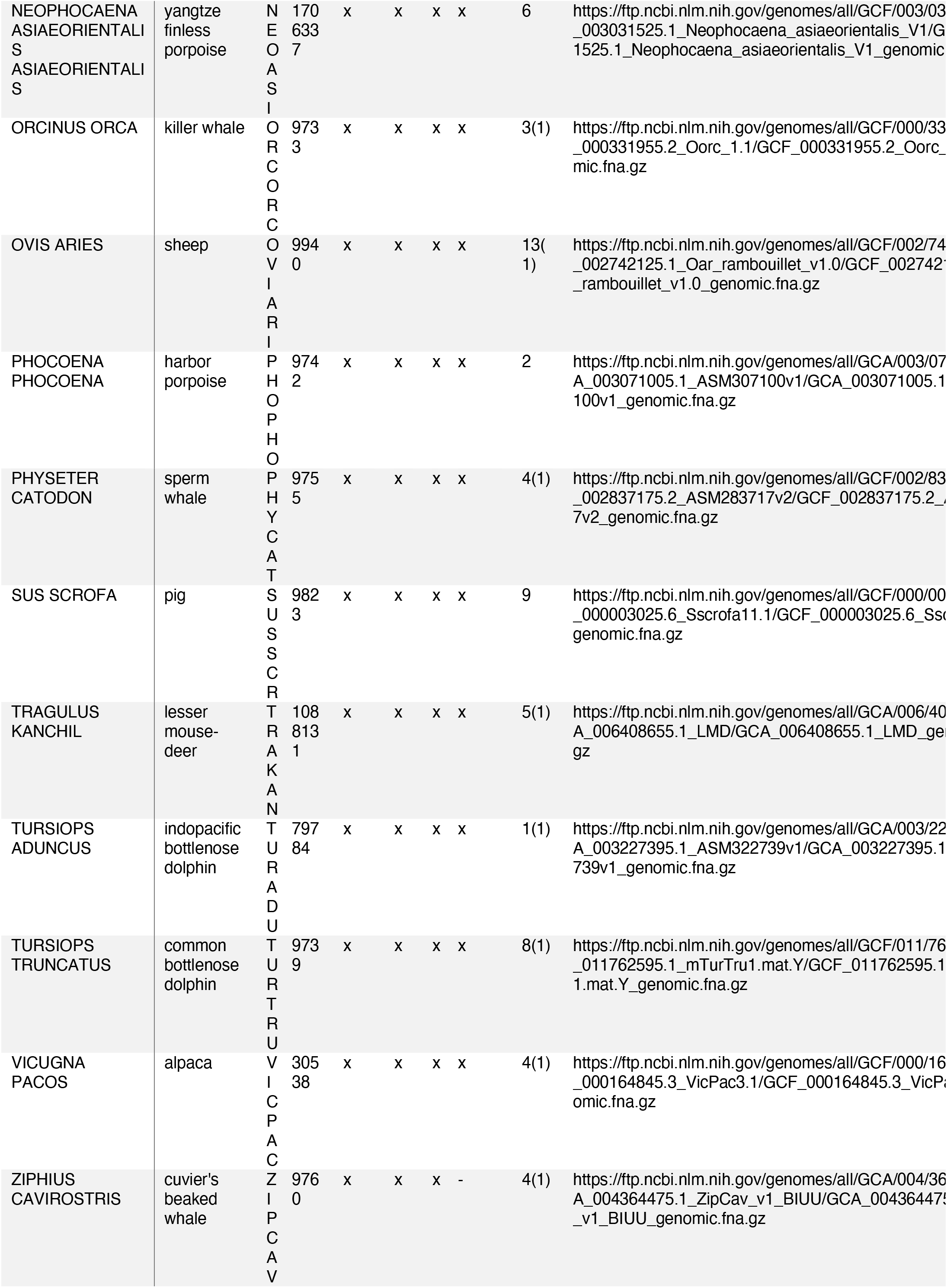
Public genome resources for PRDM9 occurrence and subsequent protein domain prediction with InterProScan. Note, that a ‘normal’ PRDM9 domain structure consists of a single C_2_H_2_ ZnF domain followed by an C_2_H_2_-ZnF-array. C_2_H_2_-ZnF prediction is based on http://zf.princeton.edu/fingerSelect.php. According to HMMER, most confident ZnF domains should have scores > 17.7, C_2_H_2_ ZnF domains with lower scores are indicated in brackets and were excluded from all downstream analyses.

**Supplementary Table 3:**
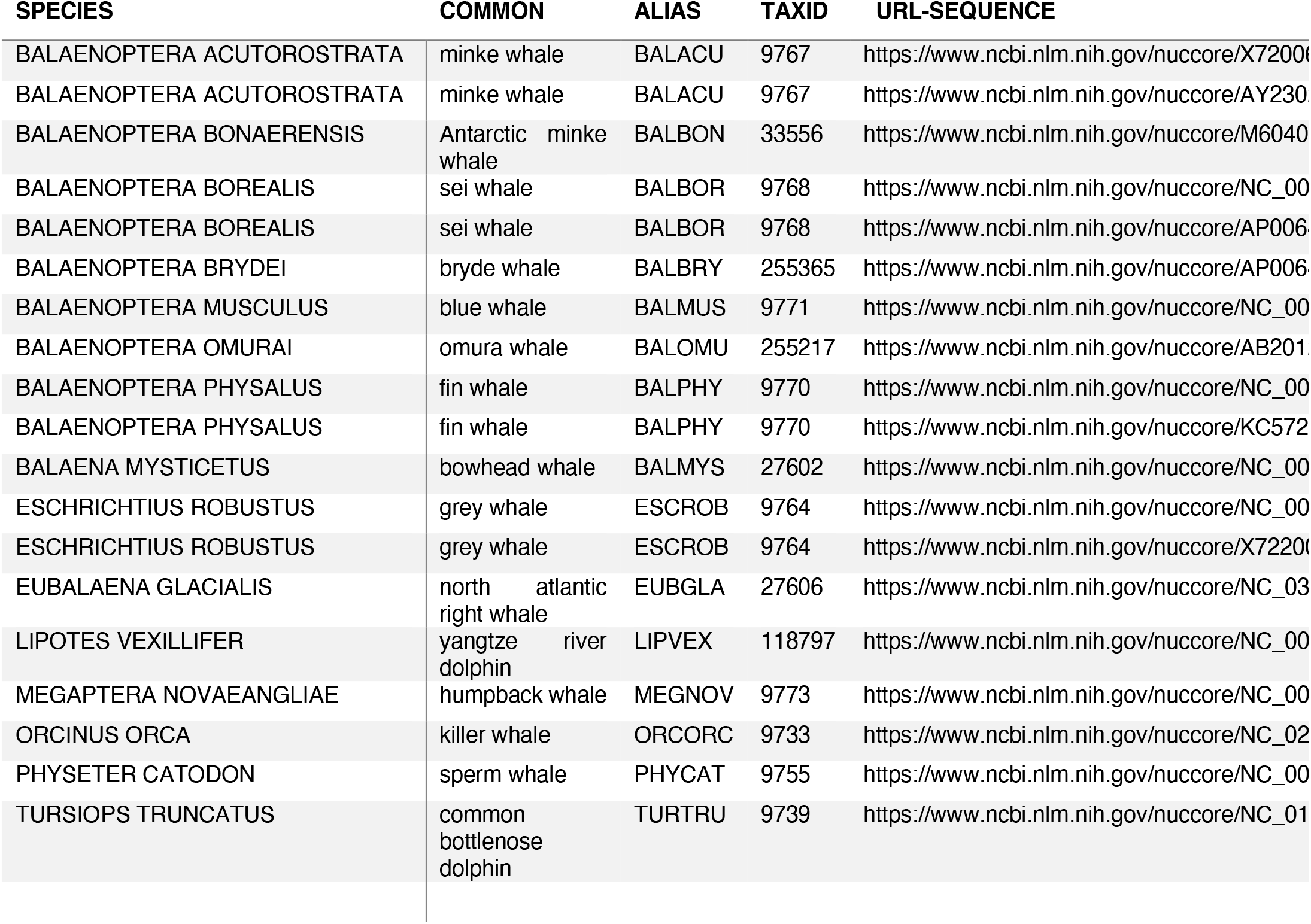
Public mitochondrial resources.

**Supplementary Table 4:**
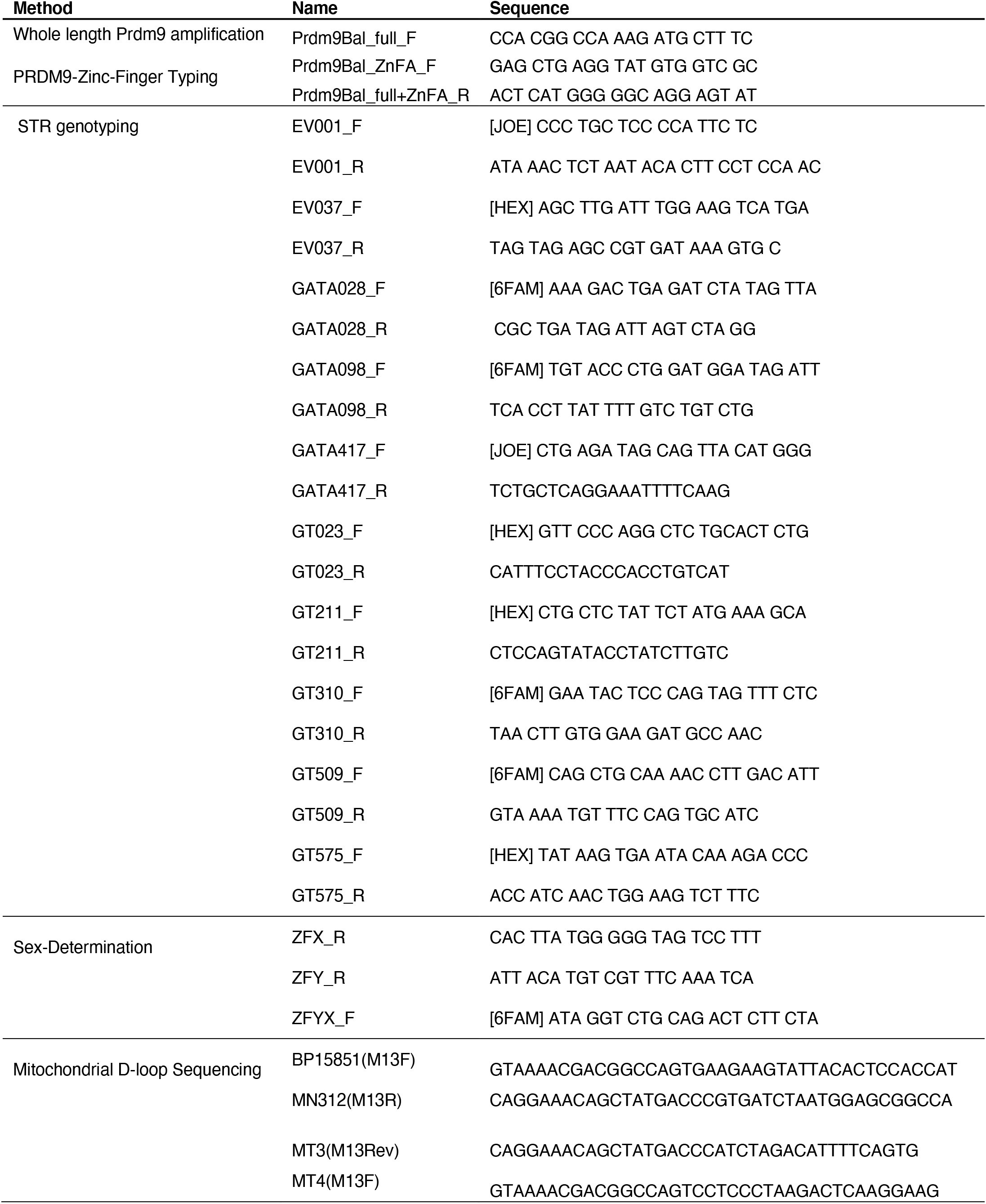
Primers used in this study.

**Supplementary Figure 6.**
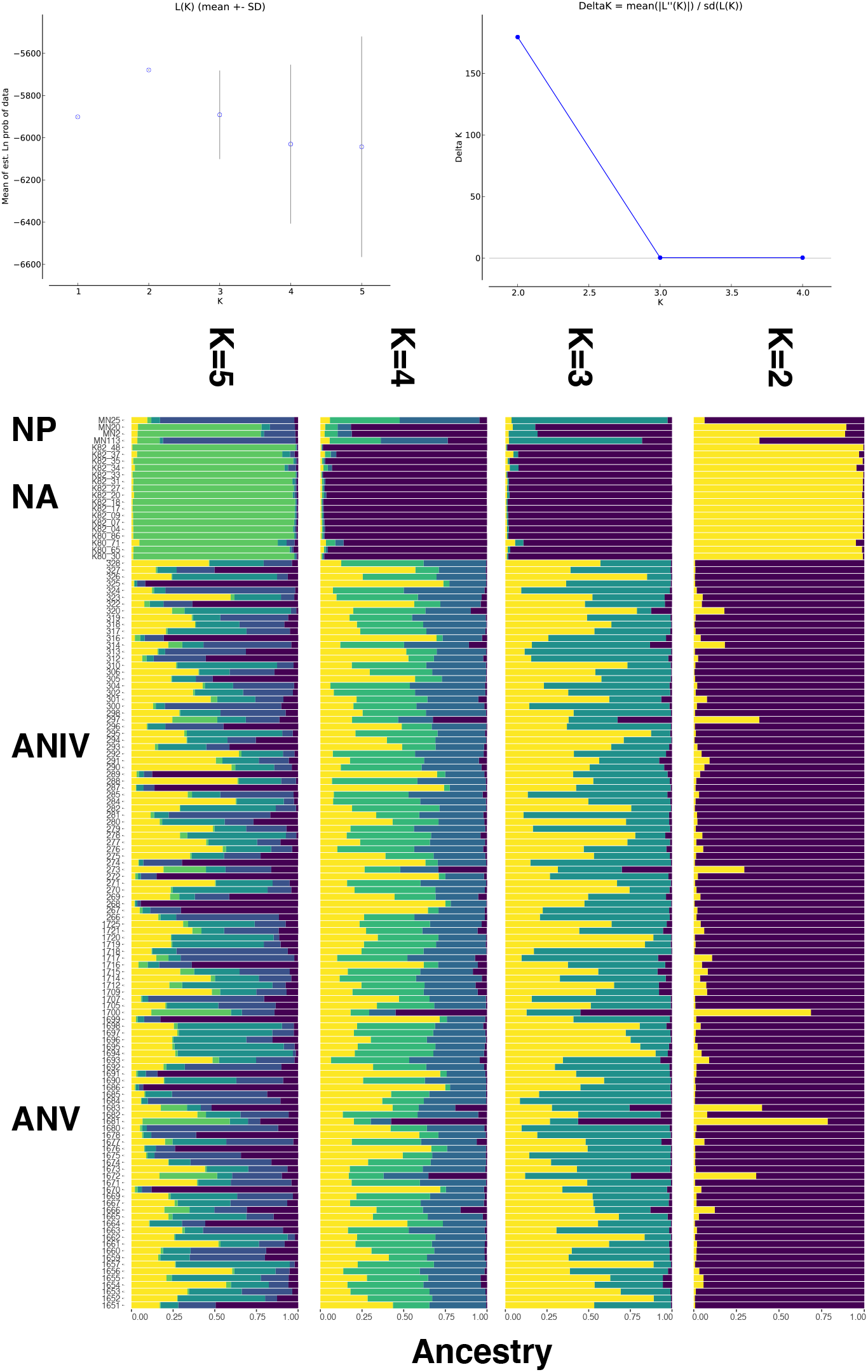
**Population structure analyses on a set of ten hypervariable microsatellite loci** using STRUCTURE without *a priori* location information (NOLOCS)

**Supplementary Figure 7.**
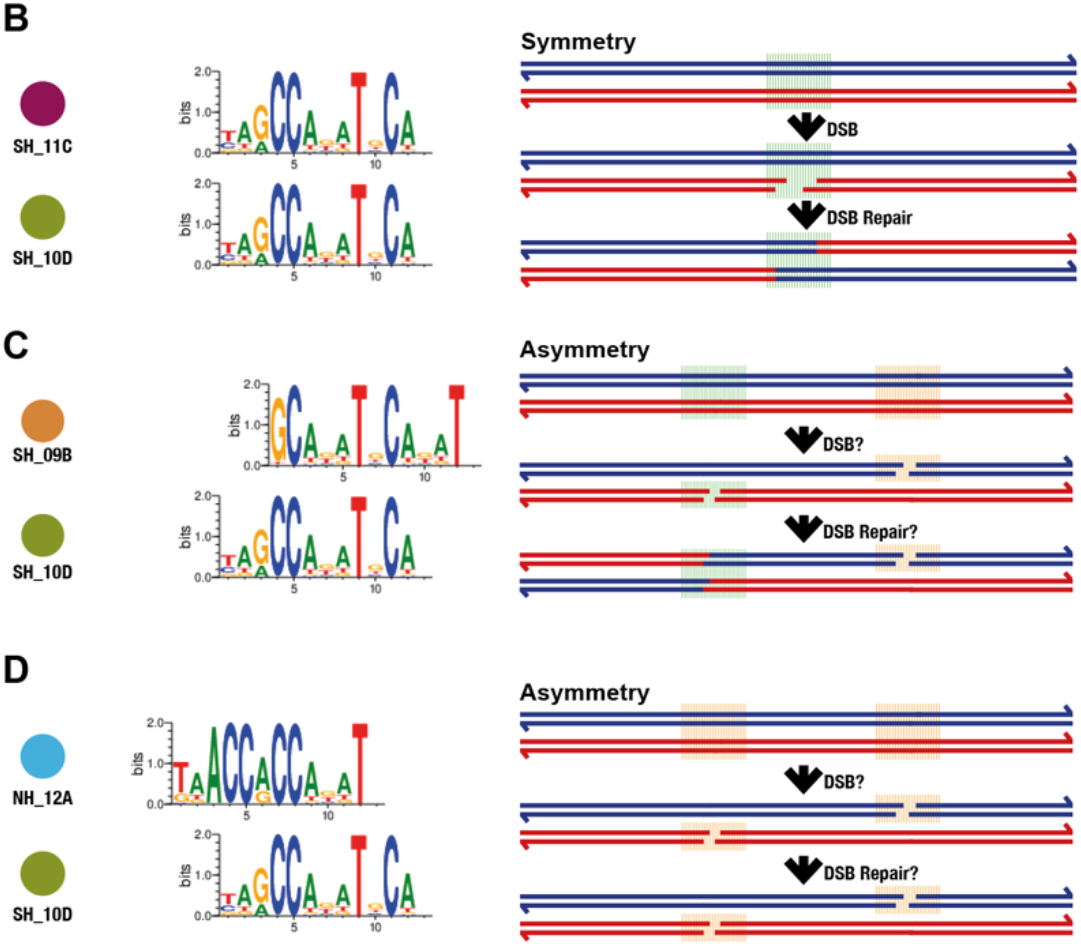
Principle of PRDM9 binding symmetry and asymmetry (A) Identical “core motif” combinations are predicted to result in fully symmetric binding and efficient DSB formation (Berg, et al. 2010; Berg, et al. 2011) (B) variants with somewhat dissimilar “core motifs”, are predicted to result in some degree of asymmetry. In humans a similar degree of motif-match still allowed DSBs necessary for successful recombination (Berg, et al. 2010; Berg, et al. 2011; Odenthal-Hesse, et al. 2014) (C) hypothetical combination of the most common variant of minke whale species, of two hemispheres, generating a putative interspecies hybrid combination. Here we would predict asymmetric positioning of recombination initiation sites, which is implicated in F_1_-hybrid male sterility in mammals (Forejt 2016).

## References

1. Imai Y, Baudat F, Taillepierre M, Stanzione M, Toth A, de Massy B: The PRDM9 KRAB domain is required for meiosis and involved in protein interactions. Chromosoma 2017, 126(6):681–695.

2. Parvanov ED, Tian H, Billings T, Saxl RL, Spruce C, Aithal R, Krejci L, Paigen K, Petkov PM: PRDM9 interactions with other proteins provide a link between recombination hotspots and the chromosomal axis in meiosis. Mol Biol Cell 2017, 28(3):488–499.

3. Berg IL, Neumann R, Lam KW, Sarbajna S, Odenthal-Hesse L, May CA, Jeffreys AJ: PRDM9 variation strongly influences recombination hot-spot activity and meiotic instability in humans. Nature genetics 2010, 42(10):859–863.

4. Berg IL, Neumann R, Sarbajna S, Odenthal-Hesse L, Butler NJ, Jeffreys AJ: Variants of the protein PRDM9 differentially regulate a set of human meiotic recombination hotspots highly active in African populations. Proceedings of the National Academy of Sciences of the United States of America 2011, 108(30):12378–12383.

5. Kono H, Tamura M, Osada N, Suzuki H, Abe K, Moriwaki K, Ohta K, Shiroishi T: Prdm9 polymorphism unveils mouse evolutionary tracks. DNA research : an international journal for rapid publication of reports on genes and genomes 2014, 21(3):315–326.

6. Buard J, Rivals E, Dunoyer de Segonzac D, Garres C, Caminade P, de Massy B, Boursot P: Diversity of Prdm9 zinc finger array in wild mice unravels new facets of the evolutionary turnover of this coding minisatellite. PloS one 2014, 9(1):e85021.

7. Schwartz JJ, Roach DJ, Thomas JH, Shendure J: Primate evolution of the recombination regulator PRDM9. Nat Commun 2014, 5:4370.

8. Steiner CC, Ryder OA: Characterization of Prdm9 in equids and sterility in mules. PloS one 2013, 8(4):e61746.

9. Ahlawat S, Sharma P, Sharma R, Arora R, Verma NK: Evidence of positive selection and concerted evolution in the rapidly evolving PRDM9 zinc finger domain in goats and sheep. 2016, 47(6):740–751.

10. Ahlawat S, De S, Sharma P, Sharma R, Arora R, Kataria RS, Datta TK, Singh RK: Evolutionary dynamics of meiotic recombination hotspots regulator PRDM9 in bovids. Mol Genet Genomics 2017, 292(1):117–131.

11. Ahlawat S, Sharma P, Sharma R, Arora R, De S: Zinc Finger Domain of the PRDM9 Gene on Chromosome 1 Exhibits High Diversity in Ruminants but Its Paralog PRDM7 Contains Multiple Disruptive Mutations. PloS one 2016, 11(5):e0156159.

12. Ahlawat S, Sharma P, Sharma R, Arora R, Verma NK, Brahma B, Mishra P, De S: Evidence of positive selection and concerted evolution in the rapidly evolving PRDM9 zinc finger domain in goats and sheep. Anim Genet 2016, 47(6):740–751.

13. Bakke I, Johansen S, Bakke Ø, El-Gewely MR: Lack of population subdivision among the minke whales (Balaenoptera acutorostrata) from Icelandic and Norwegian waters based on mitochondrial DNA sequences. Marine Biology 1996, 125(1):1–9.

14. Perrin WF, Jr. RLB: Minke Whales: Balaenoptera acutorostrata and B. bonaerensis. In: Encyclopedia of Marine Mammals (Second Edition). Edited by Perrin WF, Würsig B, Thewissen JGM, Second edn. Academic Press; 2009: 733–735.

15. Glover KA, Kanda N, Haug T, Pastene LA, Oien N, Goto M, Seliussen BB, Skaug HJ: Migration of Antarctic minke whales to the Arctic. PloS one 2010, 5(12):e15197.

16. Glover KA, Kanda N, Haug T, Pastene LA, Oien N, Seliussen BB, Sorvik AG, Skaug HJ: Hybrids between common and Antarctic minke whales are fertile and can back-cross. BMC Genet 2013, 14:25.

17. Malde K, Seliussen BB, Quintela M, Dahle G, Besnier F, Skaug HJ, Oien N, Solvang HK, Haug T, Skern-Mauritzen R et al: Whole genome resequencing reveals diagnostic markers for investigating global migration and hybridization between minke whale species. BMC genomics 2017, 18(1):76.

18. Maheshwari S, Barbash DA: The genetics of hybrid incompatibilities. Annual review of genetics 2011, 45:331–355.

19. Coyne JA, Orr HA: Speciation: Sinauer Associates, Inc.; 2004.

20. Mihola O, Trachtulec Z, Vlcek C, Schimenti JC, Forejt J: A mouse speciation gene encodes a meiotic histone H3 methyltransferase. Science 2009, 323(5912):373–375.

21. Slater GS, Birney E: Automated generation of heuristics for biological sequence comparison. BMC Bioinformatics 2005, 6:31.

22. Quevillon E, Silventoinen V, Pillai S, Harte N, Mulder N, Apweiler R, Lopez R: InterProScan: protein domains identifier. Nucleic Acids Res 2005, 33(Web Server issue):W116–120.

23. Mistry J, Finn RD, Eddy SR, Bateman A, Punta M: Challenges in homology search: HMMER3 and convergent evolution of coiled-coil regions. Nucleic Acids Res 2013, 41(12):e121.

24. Suchard MA, Redelings BD: BAli-Phy: simultaneous Bayesian inference of alignment and phylogeny. Bioinformatics 2006, 22(16):2047–2048.

25. Katoh K, Kuma K, Toh H, Miyata T: MAFFT version 5: improvement in accuracy of multiple sequence alignment. Nucleic Acids Res 2005, 33(2):511–518.

26. Nguyen LT, Schmidt HA, von Haeseler A, Minh BQ: IQ-TREE: a fast and effective stochastic algorithm for estimating maximum-likelihood phylogenies. Molecular biology and evolution 2015, 32(1):268–274.

27. Ye J, Coulouris G, Zaretskaya I, Cutcutache I, Rozen S, Madden TL: Primer-BLAST: A tool to design target-specific primers for polymerase chain reaction. BMC Bioinformatics 2012, 13:134.

28. van Pijlen IA, Amos B, Burke T: Patterns of genetic variability at individual minisatellite loci in minke whale Balaenoptera acutorostrata populations from three different oceans. Molecular biology and evolution 1995, 12(3):459–472.

29. Valsecchi E, Amos W: Microsatellite markers for the study of cetacean populations. Mol Ecol 1996, 5(1):151–156.

30. Palsboll PJ, Berube M, Larsen AH, Jorgensen H: Primers for the amplification of tri- and tetramer microsatellite loci in baleen whales. Mol Ecol 1997, 6(9):893–895.

31. Berube M, Jorgensen H, McEwing R, Palsboll PJ: Polymorphic di-nucleotide microsatellite loci isolated from the humpback whale, Megaptera novaeangliae. Mol Ecol 2000, 9(12):2181–2183.

32. Berube M, Palsboll P: Identification of sex in cetaceans by multiplexing with three ZFX and ZFY specific primers. Mol Ecol 1996, 5(2):283–287.

33. Mukaj A, Pialek J, Fotopulosova V, Morgan AP, Odenthal-Hesse L, Parvanov ED, Forejt J: Prdm9 inter-subspecific interactions in hybrid male sterility of house mouse. Molecular biology and evolution 2020.

34. Orozco-terWengel P, Corander J, Schlotterer C: Genealogical lineage sorting leads to significant, but incorrect Bayesian multilocus inference of population structure. Mol Ecol 2011, 20(6):1108–1121.

35. Earl DA, vonHoldt BM: STRUCTURE HARVESTER: a website and program for visualizing STRUCTURE output and implementing the Evanno method. Conservation Genetics Resources 2011, 4(2):359–361.

36. Evanno G, Regnaut S, Goudet J: Detecting the number of clusters of individuals using the software STRUCTURE: a simulation study. Mol Ecol 2005, 14(8):2611–2620.

37. Ramasamy RK, Ramasamy S, Bindroo BB, Naik VG: STRUCTURE PLOT: a program for drawing elegant STRUCTURE bar plots in user friendly interface. Springerplus 2014, 3:431.

38. Glover KA, Haug T, Øien N, Walløe L, Lindblom L, Seliussen BB, Skaug HJ: The Norwegian minke whale DNA register: a data base monitoring commercial harvest and trade of whale products. Fish and Fisheries 2012, 13(3):313–332.

39. Kearse M, Moir R, Wilson A, Stones-Havas S, Cheung M, Sturrock S, Buxton S, Cooper A, Markowitz S, Duran C et al: Geneious Basic: an integrated and extendable desktop software platform for the organization and analysis of sequence data. Bioinformatics 2012, 28(12):1647–1649.

40. Jeffreys AJ, Neumann R, Wilson V: Repeat unit sequence variation in minisatellites: a novel source of DNA polymorphism for studying variation and mutation by single molecule analysis. Cell 1990, 60(3):473–485.

41. Paradis E: pegas: an R package for population genetics with an integrated-modular approach. Bioinformatics 2010, 26(3):419–420.

42. Murrell B, Wertheim JO, Moola S, Weighill T, Scheffler K, Kosakovsky Pond SL: Detecting individual sites subject to episodic diversifying selection. PLoS genetics 2012, 8(7):e1002764.

43. Vara C, Capilla L, Ferretti L, Ledda A, Sanchez-Guillen RA, Gabriel SI, Albert-Lizandra G, Florit-Sabater B, Bello-Rodriguez J, Ventura J et al: PRDM9 diversity at fine geographical scale reveals contrasting evolutionary patterns and functional constraints in natural populations of house mice. Molecular biology and evolution 2019.

44. Paradis E, Claude J, Strimmer K: APE: Analyses of Phylogenetics and Evolution in R language. Bioinformatics 2004, 20(2):289–290.

45. Persikov AV, Singh M: De novo prediction of DNA-binding specificities for Cys2His2 zinc finger proteins. Nucleic Acids Res 2014, 42(1):97–108.

46. Baker Z, Schumer M, Haba Y, Bashkirova L, Holland C, Rosenthal GG, Przeworski M: Repeated losses of PRDM9-directed recombination despite the conservation of PRDM9 across vertebrates. Elife 2017, 6.

47. Paigen K, Petkov PM: PRDM9 and Its Role in Genetic Recombination. Trends Genet 2018, 34(4):291–300.

48. Myers S, Bowden R, Tumian A, Bontrop RE, Freeman C, MacFie TS, McVean G, Donnelly P: Drive against hotspot motifs in primates implicates the PRDM9 gene in meiotic recombination. Science 2010, 327(5967):876–879.

49. Jeffreys AJ, Cotton VE, Neumann R, Lam KW: Recombination regulator PRDM9 influences the instability of its own coding sequence in humans. Proceedings of the National Academy of Sciences of the United States of America 2013, 110(2):600–605.

50. Jin L, Chakraborty R: Population structure, stepwise mutations, heterozygote deficiency and thei implications in DNA forensics. Heredity 1995, 74:274–285.

51. Nei M: Analysis of Gene Diversity in Subdivided Populations. Proceedings of the National Academy of Sciences of the United States of America 1973, 70(12):pp. 3321–3323.

52. Jost L: G(ST) and its relatives do not measure differentiation. Mol Ecol 2008, 17(18):4015–4026.

53. Porras-Hurtado L, Ruiz Y, Santos C, Phillips C, Carracedo A, Lareu MV: An overview of STRUCTURE: applications, parameter settings, and supporting software. Front Genet 2013, 4:98.

54. Parvanov ED, Petkov PM, Paigen K: Prdm9 controls activation of mammalian recombination hotspots. Science 2010, 327(5967):835.

55. Cole F, Baudat F, Grey C, Keeney S, de Massy B, Jasin M: Mouse tetrad analysis provides insights into recombination mechanisms and hotspot evolutionary dynamics. Nature genetics 2014, 46(10):1072–1080.

56. Billings T, Parvanov ED, Baker CL, Walker M, Paigen K, Petkov PM: DNA binding specificities of the long zinc-finger recombination protein PRDM9. Genome biology 2013, 14(4):R35.

57. Striedner Y, Schwarz T, Welte T, Futschik A, Rant U, Tiemann-Boege I: The long zinc finger domain of PRDM9 forms a highly stable and long-lived complex with its DNA recognition sequence. Chromosome Res 2017, 25(2):155–172.

58. Groeneveld LF, Atencia R, Garriga RM, Vigilant L: High diversity at PRDM9 in chimpanzees and bonobos. PloS one 2012, 7(7):e39064.

59. Pastene LA, Goto M, Kanda N, Zerbini AN, Kerem D, Watanabe K, Bessho Y, Hasegawa M, Nielsen R, Larsen F et al: Radiation and speciation of pelagic organisms during periods of global warming: the case of the common minke whale, Balaenoptera acutorostrata. Mol Ecol 2007, 16(7):1481–1495.

60. Rosel PE, Wilcox LA, Monteiro C, Tumlin MC: First record of Antarctic minke whale, Balaenoptera bonaerensis, in the northern Gulf of Mexico. Marine Biodiversity Records 2016, 9(1).

61. Pratto F, Brick K, Khil P, Smagulova F, Petukhova GV, Camerini-Otero RD: DNA recombination. Recombination initiation maps of individual human genomes. Science 2014, 346(6211):1256442.

62. Smagulova F, Brick K, Pu Y, Camerini-Otero RD, Petukhova GV: The evolutionary turnover of recombination hot spots contributes to speciation in mice. Genes Dev 2016, 30(3):266–280.

63. Davies B, Hatton E, Altemose N, Hussin JG, Pratto F, Zhang G, Hinch AG, Moralli D, Biggs D, Diaz R et al: Re-engineering the zinc fingers of PRDM9 reverses hybrid sterility in mice. Nature 2016, 530(7589):171–176.

64. Forejt J: Genetics: Asymmetric breaks in DNA cause sterility. Nature 2016, 530(7589):167–168.

65. Zelazowski MJ, Cole F: X marks the spot: PRDM9 rescues hybrid sterility by finding hidden treasure in the genome. Nat Struct Mol Biol 2016, 23(4):267–269.

66. Dzur-Gejdosova M, Simecek P, Gregorova S, Bhattacharyya T, Forejt J: Dissecting the genetic architecture of F1 hybrid sterility in house mice. Evolution 2012, 66(11):3321–3335.

67. Bhattacharyya T, Reifova R, Gregorova S, Simecek P, Gergelits V, Mistrik M, Martincova I, Pialek J, Forejt J: X chromosome control of meiotic chromosome synapsis in mouse inter-subspecific hybrids. PLoS genetics 2014, 10(2):e1004088.

68. Tamura K, Nei M: Estimation of the number of nucleotide substitutions in the control region of mitochondrial DNA in humans and chimpanzees. Molecular biology and evolution 1993, 10(3):512–526.

